# CONSULT: Accurate contamination removal using locality-sensitive hashing

**DOI:** 10.1101/2021.03.18.436035

**Authors:** Eleonora Rachtman, Vineet Bafna, Siavash Mirarab

**Affiliations:** Bioinformatics and Systems Biology Graduate Program, UC San Diego, CA, USA; Department of Computer Science and Engineering, UC San Diego, CA, USA; Department of Electrical and Computer Engineering, UC San Diego, CA, USA

**Keywords:** Contamination filtering, *k*-mer matching, Locality-sensitive hashing, Genome skimming

## Abstract

A fundamental question appears in many bioinformatics applications: Does a sequencing read belong to a large dataset of genomes from some broad taxonomic group, even when the closest match in the set is evolutionarily divergent from the query? For example, low-coverage genome sequencing (skimming) projects either assemble the organelle genome or compute genomic distances directly from unassembled reads. Using unassembled reads needs contamination detection because samples often include reads from unintended groups of species. Similarly, assembling the organelle genome needs distinguishing organelle and nuclear reads. While k-mer-based methods have shown promise in read-matching, prior studies have shown that existing methods are insufficiently sensitive for contamination detection. Here, we introduce a new read-matching tool called CONSULT that tests whether k-mers from a query fall within a user-specified distance of the reference dataset using locality-sensitive hashing. Taking advantage of large memory machines available nowadays, CONSULT libraries accommodate tens of thousands of microbial species. Our results show that CONSULT has higher true-positive and lower false-positive rates of contamination detection than leading methods such as Kraken-II and improves distance calculation from genome skims. We also demonstrate that CONSULT can distinguish organelle reads from nuclear reads, leading to dramatic improvements in skims-based mitochondrial assemblies.

## 1 Introduction

Despite the decreased cost of whole-genome sequencing, carrying out large-scale cohort studies of non-human species using assembled genomes is still daunting (1). Low-cost sequencing projects remain an attractive alternative in biodiversity and ecological research (2, 3). Such studies can include a large number of samples sequenced at 1-2× read coverage, often called genome skims (4, 5, 6). Traditionally, genome-skimming data were used for assembling the over-represented organelle genome using one of several approaches that have been developed (7, 8, 9, 10, 11, 12, 13). More recently, noting that skimming also produces a large number of unassembled reads from the nuclear genome, researchers have been inspired to use those unassembled reads to answer questions of interest in area of biodiversity, including sample identification and population genetics (14). This vision can be realized using assembly-free and alignment-free methods where bags of unassembled reads represent *both* the labeled species (i.e., reference) and the new sample that need to be identified (i.e., query), and these bags of reads are directly compared. This vision has been pursued by several methods that enable computing distances among skims (15, 16, 17, 18, 19) and using those distances for phylogenetic placement (20, 21).

The broad use of skimming data for biodiversity is within reach, but a significant hurdle remains: contamination. Analyzing raw unassembled reads without mapping to reference genomes is particularly vulnerable to the presence of extraneous sequencing reads that do not belong to the species of interest (22, 23). Foreign DNA originating from parasites, symbionts, diet, bacteria, and human are often mixed in with supposedly single-species genome skims with the sequencing step further contributing to the contamination (24, 25, 26). With a slight abuse of terminology, we broadly refer to all external DNA outside of the genomes of interest as contaminants. Such contamination has the potential to reduce accuracy of distances estimated from genome skims. Using theoretical modeling and experimental studies, (27) have shown that contamination can lead to over and underestimation of distances between genome skims using assembly-free methods such as Skmer.

Examination of raw reads for contamination detection is not a new challenge. Early filtering techniques that relied on *k*-mer-coverage or GC content (28, 29, 30) missed contaminants frequently and were replaced by methods that use sequence similarity to search against libraries of potential contaminants (31). The current practice is to re-purpose classification methods used in the taxonomic characterization of microbial metagenomes to identify extraneous reads in genomic datasets (31). Metagenome classifiers use a variety of approaches, including read alignment as nucleotide or protein, *k*-mer mapping, and alignment of marker genes (32, 33, 34). Among these, marker-based and protein-based methods cannot be used for contamination removal as they will only detect reads from markers or coding sequence (CDS) regions. Among the remaining methods, *k*-mer-based methods (e.g., 35, 36, 37, 38, 39) are widely-used and present a reasonable compromise between speed and sensitivity. In particular, thanks to its speed, accuracy, and user-friendly implementation, Kraken-II (39) is widely used.

For contamination removal, unlike taxonomic classification, we are interested in detecting the broad taxonomic group of a read. For example, given a skim from an insect, we seek to find reads that can be rejected as belonging to Arthropoda. Thus, reads clearly belonging to prokaryotes, fungi, or plants would be judged as contamination. Metagenomic classification tools *do* classify reads at high levels but have not necessarily been optimized for higher level classification. Instead, their goal has been increasing specificity (e.g., detecting species). While detecting higher levels should be easier in principle, methods remain inadequate.

A shortcoming of metagenomic tools is their reduced ability to match reads when evolutionary close species are not available in the reference set (40, 41, 42, 43). Much of the microbial diversity on earth is not reflected with close representatives in the reference datasets (44, 45). Thus, contamination removal tools should ideally identify the broad group of species generating a read even when the reference is *sparse.* Current methods are not sensitive enough. For instance, the phylum level classification lacks sensitivity when tested on novel data (42). We recently showed (27) that even at the domain-level, the sensitivity of the leading method Kraken-II (39) degrades dramatically as the distance to the closest match in the database increases above ≈8% demonstrating that even leading metagenomic methods have serious limitations for contamination removal. The limited sensitivity of methods has spurred the development of many reference sets (46, 47, 48, 49), including recent whole-genome databases with up to 25000 genomes (50, 51, 52). However, despite their substantial size, these databases (or close to a million prokaryotic genomes available on RefSeq and GenBank databases) include only a fraction of the estimated 10^12^ extant microbial species (53). Thus, better reference datasets are not enough; we need more sensitive sequence matching methods.

Sensitive read matching tools can also help the organelle-based use of genome skims. Assembling organelle genomes needs some way of telling apart nuclear and organelle reads. Existing methods rely on either a seed-and-extend method where a seed (e.g., COI barcodes, available for millions of species) is used to find a part of the organelle genome and seek neighbouring regions in the assembly graphs (11, 12). An alternative is to rely on differential coverage of organelle and nuclear genomes to distinguish the two (7, 9). Finally, when a close reference genome is available, using that genome and read mapping can be used (e.g., 10). However, given that more than 10,000 organelle genomes from across the tree of life are already available in RefSeq, an alternative approach seems fruitful. We can build a database of all existing organelle genomes and use a sensitive read matching tool to find which reads look like they belong to the organelle genome. The assembly can then proceed simply using reads that match the database at some distance.

In this paper, we introduce a read matching method and apply it to both contamination removal and organelle read detection. Our method, called CONSULT (CONtamination Spotting Using Locality-sensitive hashing Techniques), uses *k*-mers in a query sequence to search a reference database and detects whether any of the *k*-mers match any sequence in the database allowing for inexact matches up to a user-defined threshold. The general strategy is similar to Kraken-II, except CONSULT allows mismatches using the Locality-sensitive hashing (LSH) technique; however, unlike Kraken-II, CONSULT does not currently produce taxonomic assignments. We compare CONSULT to leading methods both as a contamination removal tool and as a pre-processing step to help organelle assembly and show its superior accuracy in both settings.

## 2 Materials and Methods

### 2.1 CONSULT

#### 2.1.1 Background: LSH and the motivation to use it

Exact *k*-mer matching is not sufficient for matching reads to a database when the closest species in the reference set is evolutionary distant. Here, we always chose *k* > 20 so that *k*-mers are expected to be unique except when they derive from a common ancestor (e.g., repeats). Let us examine an example. Consider a case where the query genome is at distance *d* = 0.15 to its closest match *M* in the reference set. While due to lack of independence among adjacent *k*-mers, the probability of shared *k*-mers is hard to compute, we can still compute the expected number of *k*-mers shared between a read of length *L* = 150 and *M*, which is only (*L* – *k* + 1)(1 – *d*)^*k*^ = 0.4 for *k* = 35 (default in Kraken-II). Thus, most reads would not match the reference dataset. Existing methods have recognized the need for inexact *k*-mer matching. For example, Kraken-II masks 7 positions from each *k*-mer to increase the expected number of matches (1.3 in the previous scenario), allowing many but not all reads to match. Note that most methods avoid keeping *all k*-mers of reference genomes in the reference set, further reducing the expected number of matches per read.

We approach inexact matching using LSH, which is a widely-used hashing technique for clustering similar items or finding neighbors of a data points within a distance threshold (54). LSH uses a family of functions that hash data points into buckets so that data points near each other, and *only* such data points, are located in the same buckets with high probability. An LSH requires hashing functions that guarantee the probability of two items with distance below a desired threshold *p* falling in the same bucket is higher than that of two items with distances greater than *a* · *p* for some approximation factor *a* > 1. LSH schemes are known for many distances (55, 56, 57, 58, 59); here, we use the simplest of them: Hamming distance (HD).

Designing LSH for HD is straight-forward. We can hash a *k*-mer by simply picking a random (but fixed) position in that *k*-mer, which would put two sequences of Hamming distance *d* in the same bucket with probability 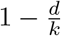. While this probability decreases with *d* (thus, forms a valid LSH), the probability reduces only linearly with d and is not expected to be effective. However, this simple hash function can be amplified using AND and OR constructions. Given two *k*-mers represented by *l* hash functions each constructed using *h* randomly-positioned bits (i.e., AND construction), the probability that *at least one* of the hash functions (i.e., OR construction) fall in the same bucket is:

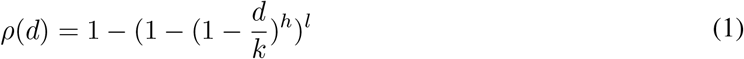

By varying *k, l*, and *h*, we can control *ρ*(*d*). Note that *l* = 1 and *h* = *k* – *s* reproduces the masking strategy used by Kraken-II (*s* is the number of masked bits). Ideally, *ρ*(*d*) should be close to 1 for *d* ≤ *p* and should quickly drop close to zero for *d* ≫ *p*. As shown in Figure S1, *ρ*(*d*) can produce an inverted S-shaped figure, and fixing *k*, many settings of *l* and *h* can lead to high *ρ*(*d*) for low distances (e.g., *d* ≤ *p* = 3) and much lower *ρ*(*d*) for higher distances (e.g., *d* > 6).

#### 2.1.2 CONSULT Algorithm

The inputs to CONSULT is set of reference genomes, represented as a set of *k*-mers, one or more query reads, and two adjustable parameter: *c* and *p*. It seeks to address the following problem: Are there at least *c k*-mers in a given read that each have at most distance *p* to some *k*-mer in the reference library? While the naive solution to this problem requires comparing each *k*-mer in each query read to each *k*-mer in the library, CONSULT uses LSH to circumvent that need. To build its library, CONSULT saves reference *k*-mers in a LSH-based lookup table (Fig. 1a), further described below. At the query time, the lookup table enables CONSULT to compare a given *k*-mer to a small (bounded) number of reference library *k*-mers to compute the HD between the query and the reference *k*-mers. A read is called a match as soon as at least c reference *k*-mers are found that match the query *k*-mer, meaning that their HD is no larger than *p* (Algorithm 1).

**Figure 1:**
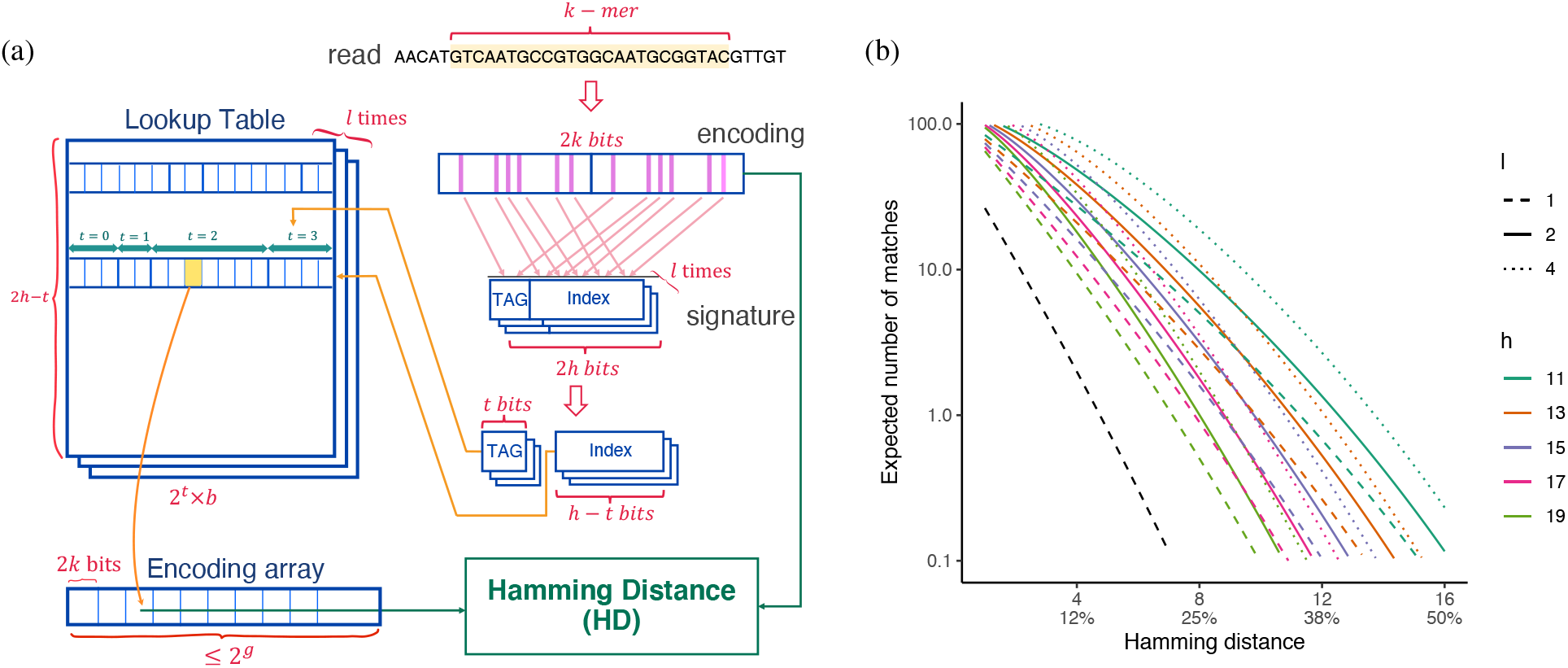
(a) A set-associative lookup table is indexed by *h* randomly-selected LSH signatures extracted from each *k*-mer and points to the encoding array. (b) We show (*L* – *k* + *1*)*ρ*(*d*): the expected number of *k*-mers that match a *k*-mer in a reference genome at distance d from the read for various *h, l* settings; *k* = 32. Black line: *k* = 35, *h* = 35 – 7 and *l* = 1, similar to default Kraken-II.

##### Encoding *k*-mers

Let’s assume we have up to 2^*g*^ *k*-mers in the reference set. Every reference *k*-mer is encoded in a 2*k*-bit number and is kept in an *encoding array* of maximum size 2^*g*^ (Fig. 1a). We use a specific Left/Right encoding that allows very fast calculation of HD using a native `popcount` instruction, an XOR, an OR, and a shift (see procedures `LeftRightEncode` and `HD` in Alg. 1).

##### Lookup Table

To find a constant-size subset of *k*-mers for computing HD, we use LSH. Hash values are generated by randomly selecting *h* 2-bits at randomly-chosen (but fixed) positions from the 2*k*-bit encodings, repeating the process *l* times to produce *l signatures.* When building the reference library, we save *l* one-to-many mappings from each *k*-mer signature to at most *b* encodings, implemented as a lookup table (Fig. 1a). When a *k*-mer is added to the encoding array, each of the *l* lookup tables is updated to point to its position, skipping a table if the corresponding row is full. The lookup tables should ideally allow around 2^*g*^ elements to accommodate all encodings, which can be achieved by setting *b* ≈ 2^*g*–2*h*^. At the query time, for each *k*-mer of a read and its reverse complement, we generate its *l* signatures (using a trick enabled by x86 instruction “extended shift” (`shld`); see `Signature` in Algorithm 2, supplementary material), which we use as index to a row of the lookup table; thus, the constant number of *k*-mers we will test equals *b* × *l*.

**Algorithm 1.**
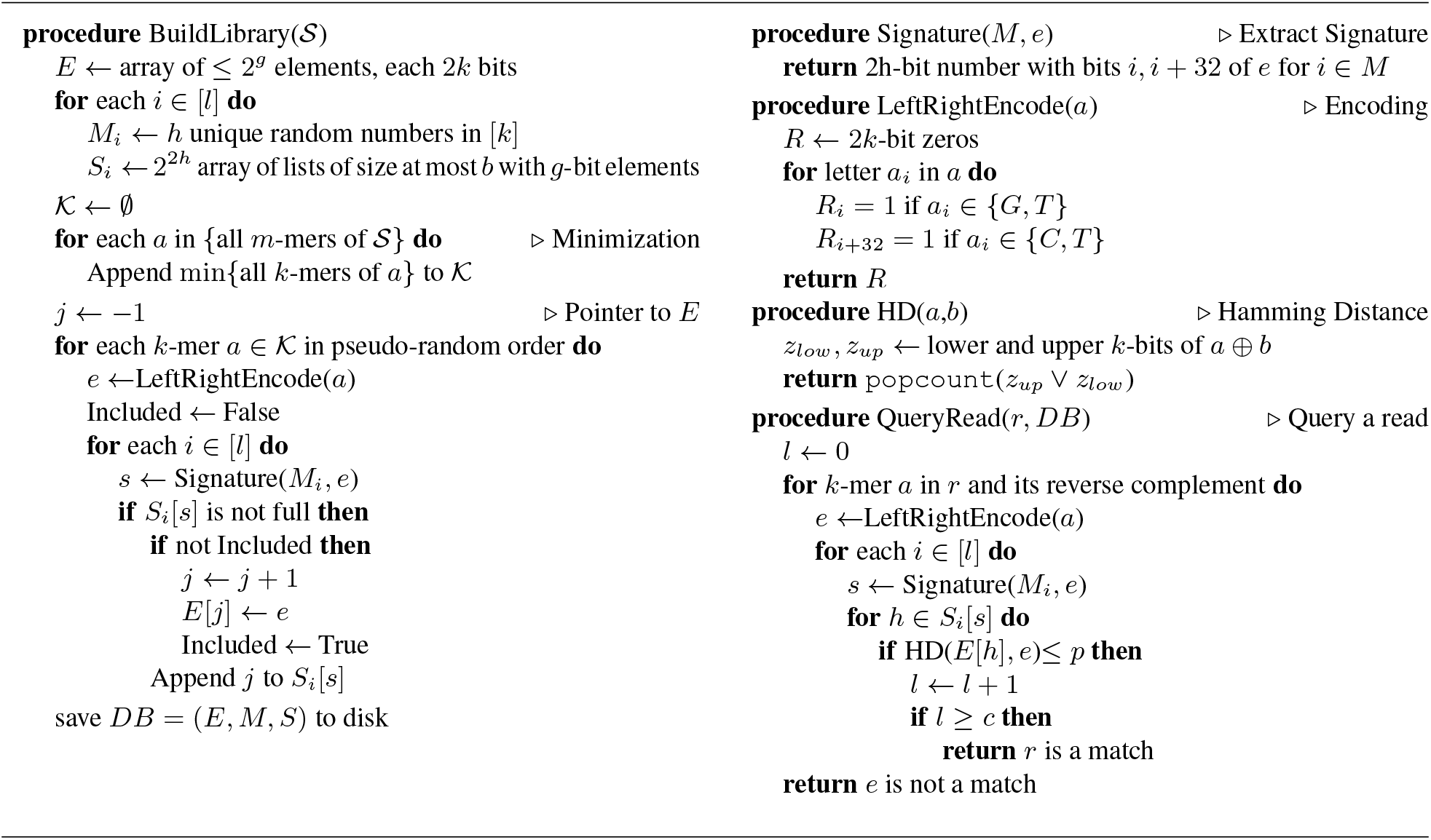
CONSULT algorithm. Here, we omit the set-associative design and tags and the fast `SHLD`-based computation of signatures for simplicity; for the more complete pseudocode, see Algorithm 2 in supplementary material. Notations: 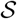: all reference sequences. Defaults: *m* = 35, *k* = 32, *h* = 15, *l* = 2, *b* = 7, *p* = 3, *c* = 1, *g* = 33. [a] denotes {0,…,*a* – 1}.

Note that there is no guarantee that signatures will appear uniformly in the reference set (Fig. S2) and so some rows can fill up sooner than others. To avoid losing *k*-mers to imbalances (Table S1), we use a set-associative lookup: The most significant *t* bits (default=2) of a signature are used as a tag and the remaining 2*h* – *t* bits as the index to the lookup table. Thus, the table has 2^2*h*–*t*^ rows, and each signature can have between 0 and *b* × 2^*t*^ entries. This design improves utilization of the table (Figs. S2, S3). We sort elements in each row by the tag and save tag boundaries.

##### HD calculation

Given that we use LSH, one may wonder why computing HD explicitly is necessary. LSH can only provide probabilistic guarantees: *k*-mers from very distant genomes have small but non-negligible chances of matching. Modern reference libraries for prokaryotes include > 10, 000 representatives genomes (51, 52), leading to 8 to 20 billion unique *k*-mers *after* minimization (Table 1). Against such huge reference libraries, small probabilities of incorrect matches blow up. To guard against false positives, CONSULT makes sure a *k*-mer is called a match *only if* its actual hamming distance to a *k*-mer in the library is below *p*. Thus, LSH is not the final arbitrator of distance; it only helps reduce hamming distance calculations. A side-effect of computing HD is that it requires keeping reference library *k*-mers in memory. For example, to keep 8 billion 32-mers in memory (our target in this study), we need 8 × 2^33^ = 64G bytes for encodings. Luckily, modern server nodes have upwards of 128GB RAM, allowing this high memory usage.

**Table 1:**
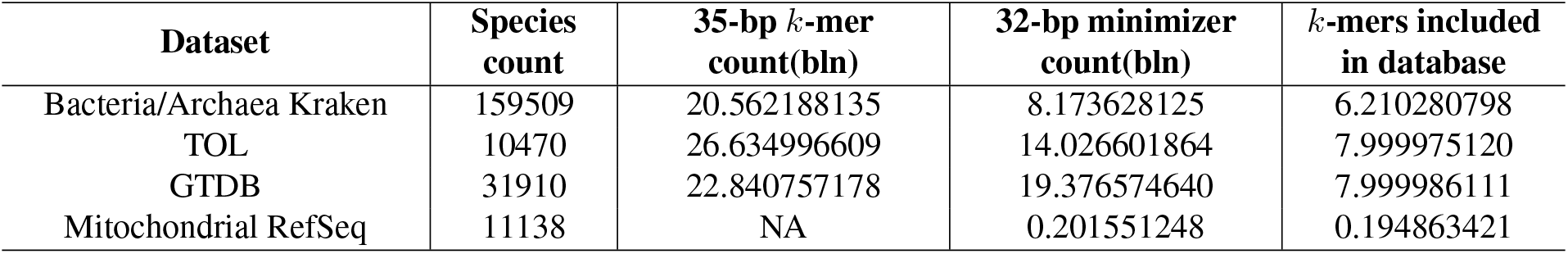
*k*-mer counts (in billions) for reference datasets before and after minimization. Number of *k*-mers corresponds to CONSULT databases constructed with default settings of the tool. Number of 35-mers indicated for Kraken before minimization computed as a sum of unique canonical *k*-mers extracted for Bacterial (19,980,385,873) and Archaeal (581,802,262) portion of the database. Subsequently Bacterial and Archaeal Kraken *k*-mer lists were concatenated and minimized. Mitochondrial *k*-mer set included all canonical 32-bp *k*-mers that were extracted without minization.

##### Parameter settings

We will explore the parameters *c* and *p* in our experiments and will select default values. The choice of *k, l*, and *h* presents intricate trade-offs between memory, running time, recall, and precision. The total memory usage is roughly 2^*g*–3^ × (2*k* + *g* × *l*) bytes. Fixing *g* = 33, to fit the entire library in a 128GB memory machine, we need 2*k* + 33 × *l* ≤ 128. Since *k* > 20 is needed for uniqueness of *k*-mers, we find *l* < 3. When the true distance of a read from a species in the database is *d*, the expected number of *k*-mers matching is *E_d_* = (*L* – *k* + 1)*ρ*(*d*). Despite dependencies, the number of *k*-mer matches is distributed around the mean. To avoid false positive matches, we want *E_d_* to be far below 1 for high distances (e.g., *d* > 40%) and to be high for small distances (e.g., *d* < 15%). As Figure 1b suggests, both *l* = 1 or *l* = 2 allow this goal in theory. The expected number of matching *k*-mers is substantially lower for *l* = 1 than *l* = 2; for example, with *k* = 32, *h* = 15, and *d* = 3, we expect 48 and 27 *k*-mers matches, respectively. Since CONSULT stops looking for matches as soon *c k*-mers match, fewer expected matches translate to longer running times. Moreover, the equation shown in Figure 1b is an over-estimate because not all *k*-mers in the reference library are in the memory. In preliminary experiments, we observed that *l* = 1 could reduce sensitivity (Table S2). Thus, we set *l* = 2 by default.

We set *k* = 32 to reduce the chances of non-homologous *k*-mer matches. As Figure 1b shows, *h* = 11 and *h* = 13 lead to many matches at a high distance, which would increase the running time. We found that *h* = 15 balances memory usage, running time, and accuracy well (Table S2). Given this choice, since our goal is to allow *g* = 33, we should ideally set *b* = 2^33–2*h*^ = 8, which unfortunately leads the library to be slightly larger than 128GB. Instead, we set *b* = 7, making our total memory usage close to 122GB for 8 billion *k*-mers (Table S2). Note that indices of the encoding array become 33-bits, but using a simple trick (keeping two encoding arrays along with an indicator bit), we can keep them as words.

##### Library construction

To build CONSULT databases, we first find all canonical 35-mers from all genomes in the reference set using Jellyfish (60) and then minimize (61) them down to 32-mers; this step reduced the *k*-mer count to include in a reference library (Table 1). We skip the minimization step for small reference datasets with < 10^30^ 32-mers. Since the Jellyfish output is pseudo-randomly ordered (62), further randomization is not needed.

### 2.2 Experimental validation

We test CONSULT in two applications: 1) as an exclusion-filtering method that seeks to find and remove contaminants among nuclear reads, and 2) as an inclusion-filtering method that seeks to detect mitochondrial reads in a genome skim to facilitate better mitochondrial assembly.

#### 2.2.1 Exclusion Filtering of contaminants

##### Reference libraries

For software validation and contamination removal testing we constructed reference libraries from three available microbial genomic datasets: Tree of Life (TOL) (52), Genome Taxonomy Database (GTDB) R05-RS95 (51) and bacterial and archaeal species present in standard Kraken-II (38, 39) (Table 1). TOL was composed of 10,575 microbial species and a reference phylogeny. Five genomes had IDs that did not exist in NCBI and were excluded from this set. The remaining genomes were assigned to the reference set (10,460 genomes), the query set (100) or both (10). GTDB included 30,238 bacterial and 1,672 archaeal genomes, that were selected to represent 194,600 samples clustered at 95% nucleotide identity. The Kraken library consisted of 158,627 Bacterial and 882 Archaeal samples available in RefSeq (as of July 2019). Both Kraken and GTDB reference sets were used without modification.

##### Experiments

We performed three experiments to test exclusion filtering of contaminants.

###### (i) Controlled distances

Similar to (27), we first evaluated the ability of CONSULT to find a match when the query is within a range of phylogenetic distances to the closest species present in a database. To control the proximity of the query to its closest match in the reference library, we selected 100 genomes from TOL such that their distances to their closest species in the tree uniformly covered a broad range of [0.0-0.3). These queries were removed from the reference set and remaining TOL genomes were used to construct CONSULT database. We also randomly selected 10 genomes to keep in both the query set and reference set which allowed us to evaluate TP and FN. Subsequently, all queries were divided into bins based on their distances to the closest match in a reference database (Table S3) and 10 plant genomes (Table S4) were added to the set of queries in every bin. Plant species are from a different domain of life compared to the TOL reference set and should not match the library; thus, they allowed us to measure FP and TN. All distance values in this experiment were computed using Mash (63). Reads for the TOL query set were simulated at 10MB using ART (64) (see Appendix C).

###### (ii) Novel genomes

We next assessed the ability of CONSULT to match genomic reads that belonged to novel microorganisms not observed in reference sets. To generate queries we used samples from Global Ocean Reference Genomes (GORG) that is a collection of 12,715 marine Bacterial and Archaeal single-cell assembled organisms (42). Marine microbial species are known to be poorly represented in public repositories (65). Since very few reads from these samples are expected to map to the reference genomes, they represent a particularly challenging classification case for databases with standard compositions and were a suitable set to test in this experiment. To generate queries, we obtained GORG assemblies from NCBI (project PRJEB33281; five assemblies were missing) and simulated query reads for every sample at 1x coverage using ART with the same settings as TOL (see Appendix C). We also included the same 10 plant species as TOL to compute the FP rates.

###### (iii) Real skims

We tested ability of CONSULT to remove contaminants from real genomic sequencing reads. For studying real genome skims, we obtained high-coverage raw SRA’s of 14 Drosophila species (Table S5) from NCBI (PRJNA427774) and subsampled them down to 200 MB bp using seqtk (66). We removed adapters, deduplicated these samples and merged paired-end reads using BBTools (67) (see Appendix C). To filter out human reads, we queried Drosophila samples against the Kraken database that included only the human reference genome, and subsequently extracted unclassified reads to use in contamination removal experiment. Drosophila reference assemblies available from (68) were used to compute true distances.

##### Tools compared

We compared performance of CONSULT with Kraken-II (38, 39), CLARK (37), CLARK-S (36) and Bowtie2 (69, 70). These are among leading identification tools based on recent benchmarking studies (71, 72, 73, 74). Kraken-II is a taxonomic sequence classifier that maps *k*-mers of the query to the lowest common ancestor (LCA) of all genomes known to contain a given *k*-mer. We constructed Kraken reference libraries for genomes that belonged to TOL, GTDB, and Bacterial/Archaeal portion of the standard Kraken database. Kraken reference libraries were built without masking low-complexity sequences, but using default settings otherwise. We note that (27) found defaults were the most effective setting for contamination removal.

CLARK, CLARK-S, and Bowtie2 are tested only in the experiment (i), and thus, their reference databases were built using the TOL dataset. CLARK is a method that does supervised sequence classification based on discriminative *k*-mers. We constructed the CLARK database using standard parameters (e.g., *k*=31 default classification mode). We set taxonomy rank to phylum (default is species) to achieve better sensitivity for contamination removal (see Appendix C for details). CLARK-S is a CLARK version that exploits multiple spaced *k*-mers and offers higher sensitivity at the expense of more RAM and slower classification speed. CLARK-S database was constructed on top of the custom CLARK database described above and querying was performed using default full mode of classification (see Appendix C). Bowtie2 is a standard general-purpose alignment tool. We built the Bowtie index reference for TOL genomes and performed local alignment of the queries using highest sensitivity setting (see Appendix C).

##### Evaluation

In experiments (*i*) and (*ii*) we report the recall and false positive rate: matched (unmatched) prokaryotic reads are TPs (FNs) and matched (unmatched) plant read are FPs (TNs). On the TOL dataset, we also compared running time and memory consumption of all tools for running a randomly sampled set of 30 small 10Mb queries from the TOL query set (Table S7). To reduce the impacts of database loading on running time, we report results when the query is a single file concatenating 15 Drosophila skims sampled at 2G bp (Table S8). In experiment (*ii*), we also report the percentage of reads from each microbial genome that match.

In experiment (*iii*), based on the results of the first two experiments, Drosophila genome skims were filtered against GTDB database, and Skmer was used to compute distances between all pairs of samples before and after filtering. Distance values obtained from Drosophila assemblies were considered the ground truth. We computed relative distance error for every sample before and after filtering in order to identify whether contamination removal improved distance estimates.

#### 2.2.2 Inclusion-filtering of mitochondrial reads to help organelle assembly

Here, we test whether CONSULT can help improve the quality of mitochondrial assembly by finding mitochondrial reads in a genome skim without a need for the standard seed-and-extend approach, the use of a very close reference genome, or reliance on coverage differences. To do so, we constructed a CONSULT reference database out of all 11,138 mitochondrial genomes available in NCBI (RefSeq release 204, January 04, 2021), which included ≈ 200 million 32-mers (Table 1). We then asked whether CONSULT can use this broadly sampled database to identify mitochondrial reads in SRA files, including from species not present in the reference set.

We base our experiment on data from the DNAMark project that skimmed 210 vertebrate species (NCBI project PRJNA607895) and attempted to assemble their mitochondrial genomes (75). We selected 42 (Table S6) out of 210 samples as follows. We included all 18 samples where the pipeline used by (75) failed to assemble mitochondrial genomes, all 6 samples that produced poor quality short contig (3–10.5 kbp) assemblies, and 18 randomly selected *good* samples with contig length >12 kbp, used as a positive control. Our SRAs include 24 species not present in the CONSULT reference dataset and 14 species not represented at the genus level (Table S6).

We compare three assembly pipelines. First, we include assemblies deposited by (75) generated using Novoplasty (11). For each of the 42 samples, we preprocessed the raw SRA files by removing adapters using AdapterRemoval (76) and merging paired-end reads with BBTools (67) (see Appendix C). We the assembled these *unfiltered* reads using plasmidSPAdes (9), which relies on read coverage to distinguish nuclear and organelle genomes. In the third approach, we first used CONSULT to searched preprocessed reads against the reference mitochondrial database and then used only the matching reads as input to SPAdes with default settings (77) to obtain the assembly.

To assess the completeness of the assemblies, we first annotated all three assemblies (original, unfiltered, and filtered) using MITOS (78) to find the known mitochondrial genes. We report the total length of the largest mitochondrial contig, gene counts for different gene groups (protein coding genes (PCG), rRNA, tRNA), and identities of annotated genes for PCG and rRNA. The length of mitochondrial genomes should be approximately ~16 kbp in size and the number of genes should be close to 37 (79).

Finally, note that we select one contig as the final mitochondrial assembly for each of the three methods. For original assemblies, only a single contig is available. For new assemblies, we mostly take the the longest contig (as long as it is ≥200 bp) as the assembly. However, in assemblies produced from unfiltered reads, the largest contigs sometimes had nuclear origin. In such cases, we instead use the longest annotated contig with at least one annotated mitochondrial PCG or rRNA gene. The identity of mitochondrial contigs was additionally verified by MitoZ (80). If no PCG or rRNA genes were assigned to any contigs in assembly, the generated reference is considered as having failed annotation.

## 3 Results

### 3.1 Exclusion filtering of contamination from nuclear reads

#### (i) Controlled distances

In the controlled distance experiment, CONSULT has the best recall among the methods that are able to control the FP rate (Fig. 2a). CLARK-S, which is specifically designed to match species absent from a reference database, has the highest sensitivity but FP rates close to 62%, making it ineffective for contamination removal. CLARK has very low FP rates (0.5%) but also much lower recall than other methods. Overall, Bowtie has a similar recall to Kraken-II with a substantially lower FP rate (1.2% versus 4.9%). CONSULT is slightly better than Kraken-II in terms of FP (4.3%) but improves recall over Kraken-II and all other tools substantially. All the tools are able to match almost all prokaryotic reads to the database when the query has an exact match in the database, and all tools have at least 91% recall when the closest match in the reference library is up to 5% distant from the query. Substantial differences between methods appear when the closest match is > 5% distant to its closest match. For example, for queries at 5–15% distance to the reference set, CONSULT matches 78% of reads while Bowtie and Kraken-II match 66% and 61%, respectively.

**Figure 2:**
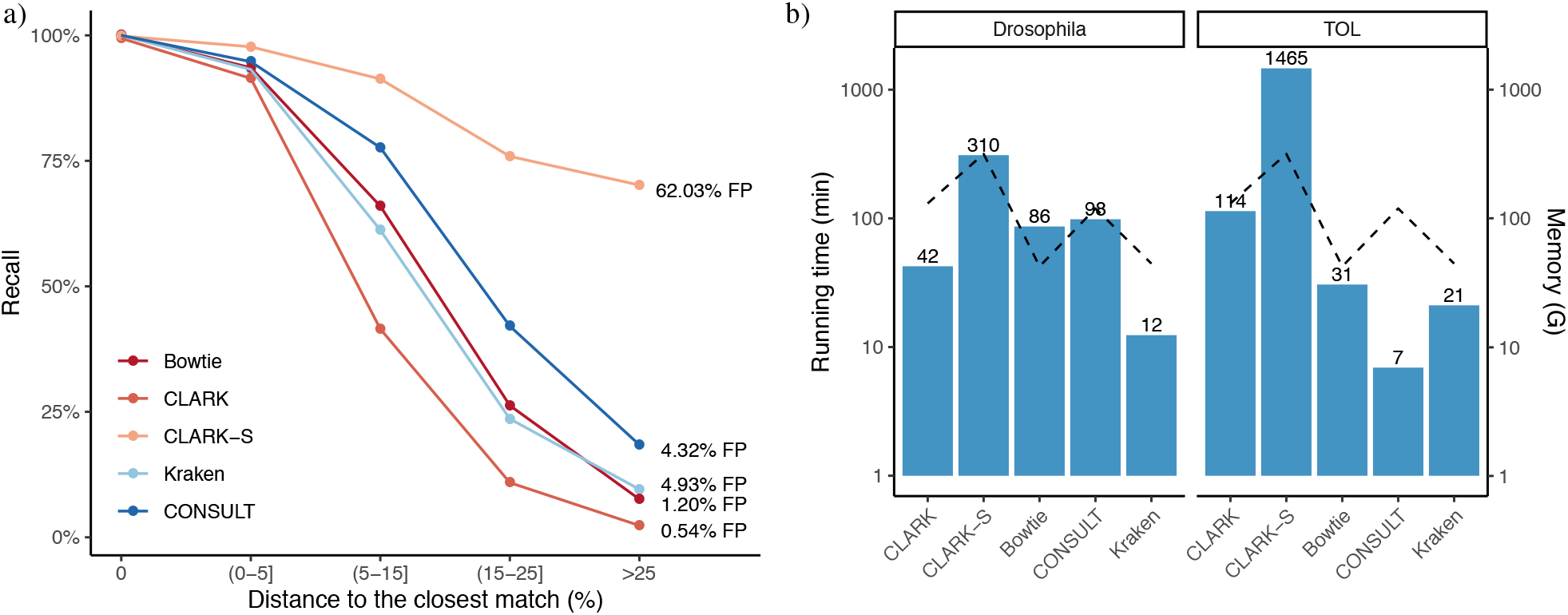
Results on the simulated data. (a) Lines show recall for different query bins for five tools run in default settings (see Methods). Each line is labeled with its associated FPR, computed using plant genomes added to the query set, which are identical across bins; thus, FPR is identical in all bins. (b) Processing speed and memory consumption for different tools searched against the TOL library using the same settings as part (a). Drosophila benchmark set included a single 30G bp Drosophila query. TOL set was composed of 30 10MB microbial queries. Computation on a machine with Intel Xeon 2.2 GHz CPU using 24 threads and 350G of RAM.

The running time of CONSULT on the TOL DB is comparable to Bowtie but slower than CLARK and Kraken-II when tested on large query files (Fig. 2b). With multiple small query files, while CONSULT is the fastest, timing is hard to interpret because CONSULT analyzes all inputs in a run, amortizing the DB load time, while others need to be run per query file (a simple issue to fix). Bowtie and Kraken-II have the lowest memory footprint, followed by CONSULT, which uses 120GB.

#### (ii) Novel genomes

Next, we turn to the GORG dataset, where CONSULT matches a far larger number of microbial reads to the reference libraries compared to Kraken-II, regardless of the reference database (Fig. 3a). CONSULT matched more reads than Kraken-II for 88% of the microbial species when searched against GTDB (Fig. S4). CONSULT and Kraken-II match at least 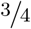 of reads for 61% and 43% of genomes, respectively. Comparing the three databases, GTDB results in the most matches for both methods, followed by TOL and Kraken. Not only does CONSULT match more reads (has higher recall), it has fewer false positives, especially for GTDB (Fig. 3b). In default settings, CONSULT controls the FP rate at 7.0% on the large GTDB dataset, whereas Kraken-II has 11.5% FP. Kraken takes around 3 minutes to load the GTDB database (on a machine with 350GB of RAM), which is substantially larger than the load time of TOL database (half a minute).

**Figure 3:**
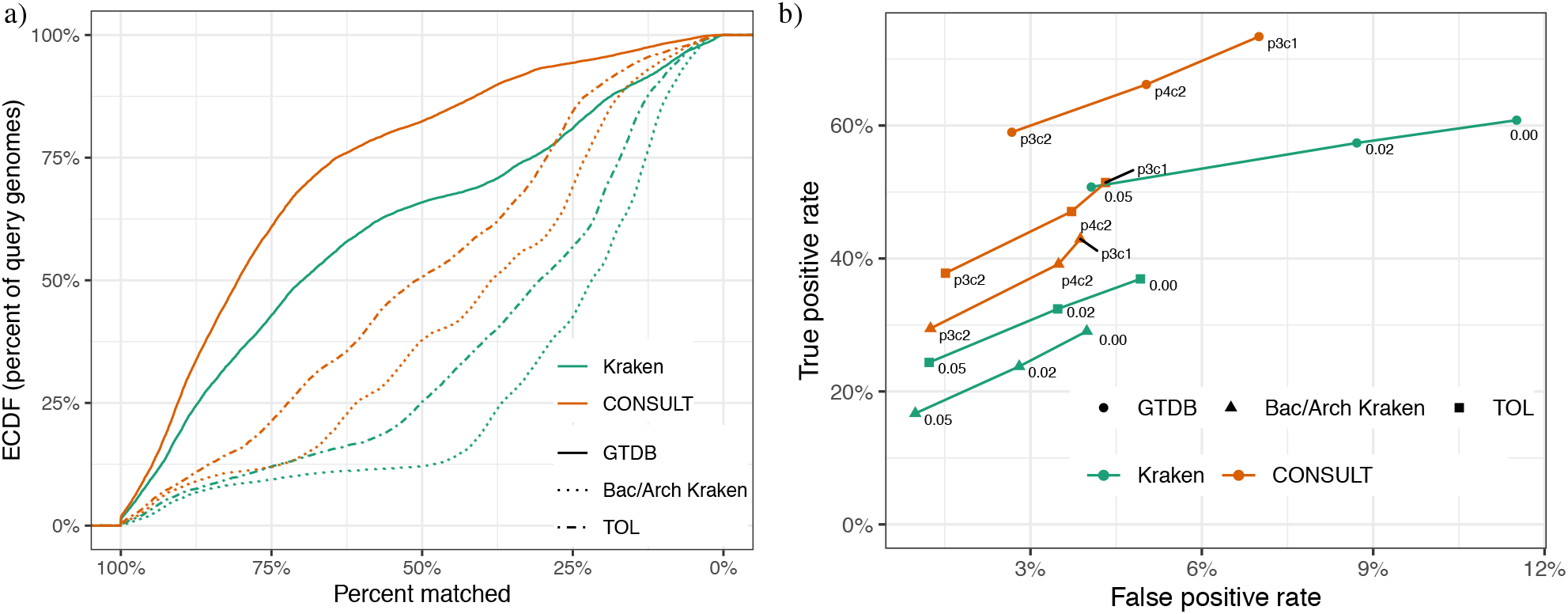
CONSULT vs Kraken-II on GORG dataset. (a) The empirical cumulative distribution of the percentage of reads in each microbial GORG genome matched to each reference database (GTDB, TOL, B/A Kraken); a point (*x*%, *y*%) means for *y*% of GORG genomes, ≥ *x*% of the reads matched the DB. (b) The ROC curve showing the mean of recall vs FP rate (i.e., plant queries matched to a DB) for both tools with three settings. Kraken-II was run with confidence level ∈ {0.00, 0.02, 0.05}. CONSULT libraries were searched using *p* ∈ {3, 4, 5, 6} and *c* ∈ {1, 2, 3, 4}; see Figure S5 for all combinations. Here, we show three low FP settings.

We next tested more strict versions of both methods (Figs. 3b). We reduced false positive rates for CONSULT by changing the (*c, p*) settings and for Kraken-II by increasing its *α* (i.e., percentage of *k*-mers in query sequence required for classification). For all levels of FP rate, CONSULT had better recall than Kraken-II for all databases tested. Thus, CONSULT performs better than Kraken. Adjusting the (*c, p*) setting of CONSULT trades off recall and FP rate (Fig. S5). For example, allowing up to 4 mismatches between *k*-mers in query and reference library produces more liberal (*c* = 1) or more conservative (*c* = 2) settings compared to the default where 3 mismatches are allowed. These combinations of parameters might be recommended for situations where a stricter FP control is required (*c* = 2) or when FP is less damaging (*c* = 1). All *p* ≥ 5 and *c* ≤ 2 lead to very high FP; e.g., *p* = 6, *c* = 1 leads to 100% recall but also 90% FP rate (Fig. S5).

#### (iii) Real skims

Testing CONSULT on real genome skims from Drosophila demonstrates that Skmer distance calculation can improve dramatically as a result of filtering (Fig. 4). Errors are reduced by as much as 44% between pairs of species (Fig. 4b). While distances tend to be underestimated before filtering, they tend to be slightly overestimated after filtering (Fig. 4a). CONSULT removes between 3.9% and 10.2% of reads from these Drosophila genomes, which is slightly lower than what Kraken removes (8–14%). These differences are consistent with the 4% higher false positive rate of Kraken-II in our simulations, suggesting that Kraken-II may be over-filtering. Overall, error after CONSULT filtering is slightly less than Kraken-II filtering in most cases (Fig. S6); these differences, while small, are statistically significant (two-way t-test *p*–value≪ 10^−4^).

**Figure 4:**
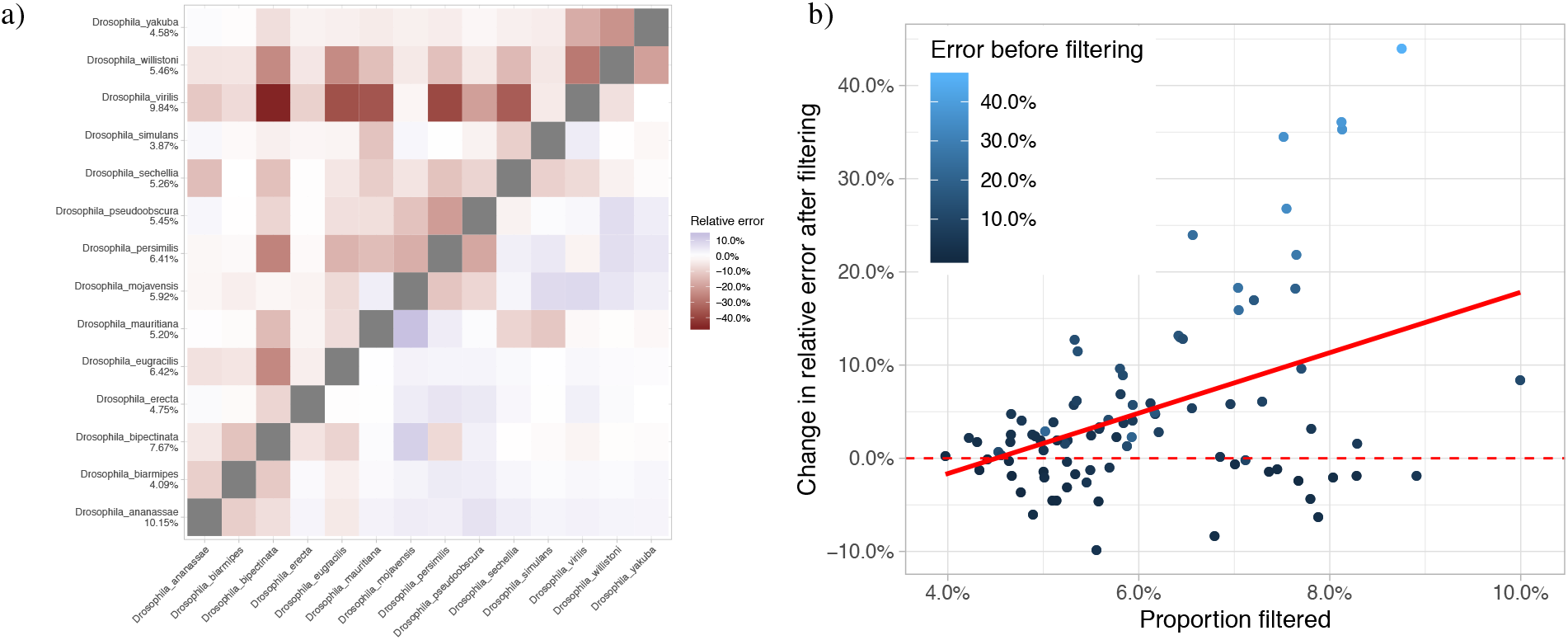
CONSULT on Drosophila skimming data. (a) Relative distance error before (upper triangle) and after (lower triangle) filtering per pair of Drosophila species. Numeric labels on y-axis represent percentage of bacterial/archaeal reads filtered per sample. (b) Change in the distance error after filtering compared to the error before filtering versus amount filtered (mean of both species); positive values indicate reduction in error. Each dot represents a pair of species.

### 3.2 Inclusion-filtering of organelle reads

Using CONSULT to find organelle reads before assembly dramatically improves the quality of the assembly, both compared to the unfiltered approach that relies on coverage and seed-and-extend method used in the original study (Fig. 5).

**Figure 5:**
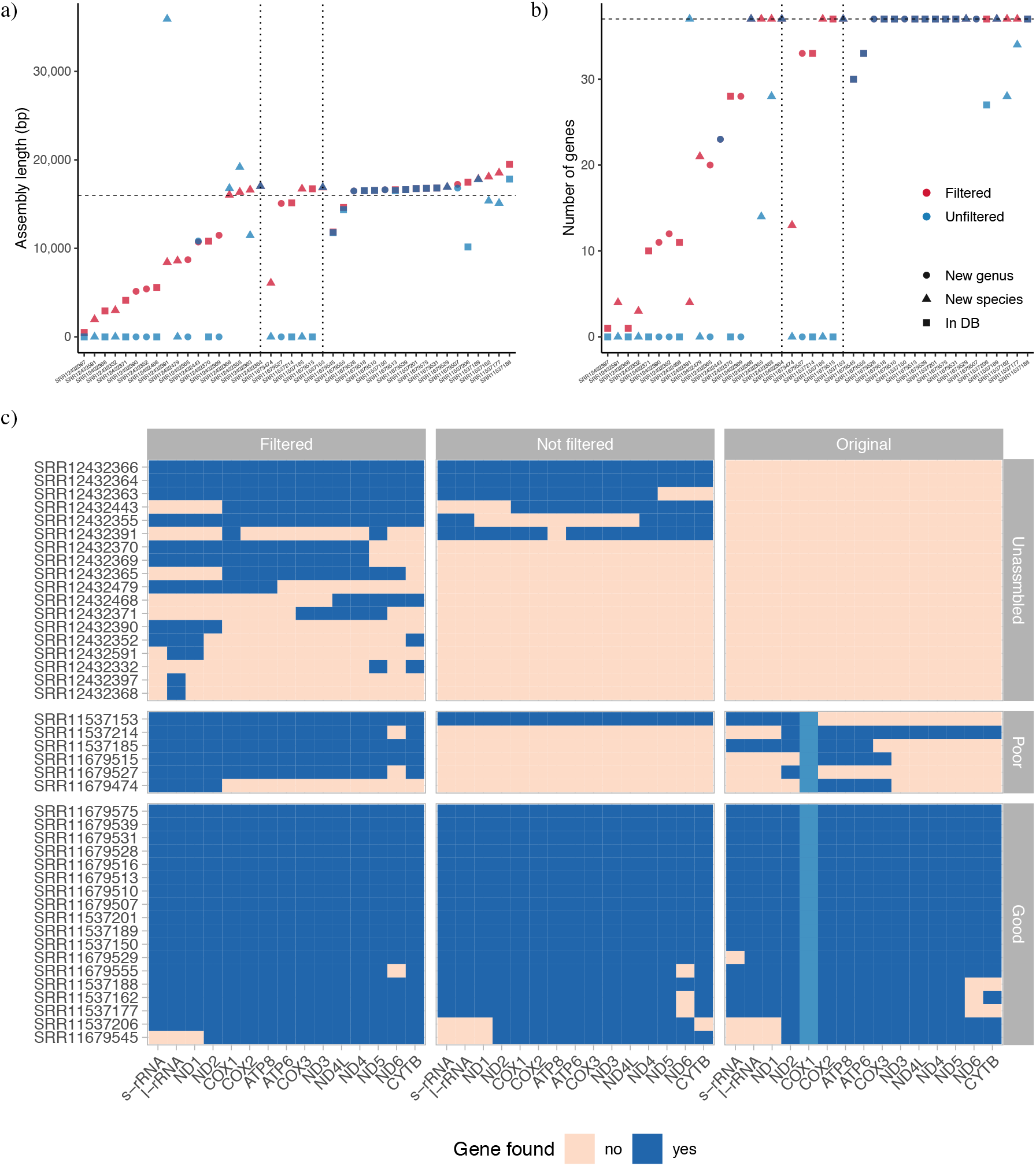
Mitochondrial assembly results. (a) Comparison of contig length between filtered and unfiltered samples. Dot shapes indicate whether any assembly from the same species or the same genus was a part of the RefSeq reference set that was used to construct CONSULT database. From left to right: sample that belonged to previously unassembled, poorly assembled, or good quality assemblies (boundaries indicated by vertical dots). The horizontal line is at expected lengths: 16 Kbp. (b) Total mitochondrial gene count for filtered and not filtered samples (counting unique PCG, rRNA and tRNA genes). Vertical line at expected number of genes: 37. (c) Mitochondrial genes identified in assemblies for filtered, unfiltered and original references. Light blue color highlights COI (COX1) gene that was used as a seed sequence for assemblies generated by the DNAMark project. In the unfiltered case, three samples (SRR12432370, SRR12432371 and SRR12432397) did not generate any contigs while 14 others generated contigs but did not have any genes annotated.

When using raw reads we obtained complete or partial mitochondrial assemblies for 25 out of 39 assembled samples. Three samples didn’t generate any contigs. Remaining 14 samples produced contigs of variable size but failed in annotation since estimated length of PCG and rRNA genes appeared shorter than expected. In contrast, when preceded by CONSULT filtering, reads were successfully assembled for all 42 samples, including the 18 samples that were left unassembled in the original study and 6 that had poor assemblies.

Assemblies produced by filtered reads were in all but one case either longer or comparable in size in comparison to assemblies generated by unfiltered reads (Fig. 5a). Similarly, they had a higher number of mitochondrial genes annotated in all but one case (Fig. 5b). The exception is the sample SRR12432391 that leads to a 35,920 bp contig when assembled from raw reads. This length is almost twice the average length of the mitochondrial genomes, which indicates a possible mis-assembly or chimeric contig. After filtering, 29 of the 42 samples had at least 27 out of 37 genes and 12 out of 15 non-tRNA genes annotated.

The completeness of an assembly after CONSULT filtering was not impacted by the presence of the corresponding species or genus in the RefSeq reference set (Fig. 5a, b). Cases where assemblies after filtering remain incomplete include both novel and observed samples. For example, of the 13 assemblies with less than 12 non-tRNA genes, four were represented exactly in the database, five had genomes from the same genus and four were not present at either level. Among the nine samples with no representation from the same genus, in five cases, CONSULT filtering improved gene recovery between 11 to 33 genes, made no difference in one case with an incomplete assembly, and recovered full assemblies in three remaining cases. Thus, effective filtering did not require representation from the same species or genus in the CONSULT reference library.

Comparison of gene annotations between newly generated and original assemblies (Fig. 5c) demonstrated that filtering enabled successful assembly for the most challenging low coverage samples. Thus, for samples that failed in the original study, we generated complete or nearly complete assemblies with up to 10 PCG identified and contig length ≥ 8719 bp for eight samples. Six samples produced partial assemblies and only four samples had contig length ≤ 3003 bp; even for them, we had some mitochondrial genes identified. For poorly preserved samples, we generated near-complete references for five out of six samples. In one case (SRR11679474), the main contig had only three PCGs and rRNAs genes, but even this sample contained all remaining genes scattered across five smaller contigs that the assembler did not merge with the longest contig. Unsurprisingly, the original seed-and-extend approach is biased toward the region including COX1, which is the seed, whereas filtered and unfiltered assemblies show no such bias. For the set of good quality samples, filtering improved gene recovery in three cases compared to unfiltered ones and five samples compared to the original assemblies; only one gene was recovered by one of original assemblies but not the filtered ones.

Additionally, since filtering reduced the number of sequencing reads that are being assembled, we observed ≈ 7× running time improvement with filtered versus unfiltered reads (estimate includes CONSULT time), going from ≈ 65 min to ≈ 9.7 min per sample. This speed-up was calculated for 24 SRR that belonged to poor and good assembly groups (120G of RAM, 24 threads).

## 4 Discussion

We introduced CONSULT, a general purpose *k*-mer based read matching tool that might help in a variety of applications where there is a need to separate sequencing reads of interest from extraneous reads outside of the group. By careful engineering of the software, we have made it possible to run CONSULT on large reference datasets (e.g., tens of thousands of prokaryotic species). Our results on contamination removal showed that when the closest species in the reference set was substantially distant (≈15–20%) from the query, CONSULT improved upon existing methods such as Kraken, CLARK(-S), and Bowtie2 both in terms of sensitivity and specificity. Our results on assembling organelles from genome skims showed that using CONSULT to pre-filter reads that seem to belong an organelle genome can improve assembly quality and speed.

CONSULT is based on LSH, a concept that has been used before for sequence analysis (81, 82, 83, 84, 85). Unlike earlier methods, CONSULT takes advantage of the large memory available on modern machines, which were traditionally not available. Nowadays, we can easily afford to keep billions of 32-mers in memory; thus, we can use LSH only to find a small set of *k*-mers for which we compute distances exactly. As such, CONSULT does not have false positives (in the sense that it guarantees every match is below the desired threshold). It only can have false negatives. However, missing some *k*-mer matches is tolerable because if a read truly belongs, its other *k*-mers must match. Consistent with this observation, our data show that even when we include half or one-third of *k*-mers from a reference dataset in the memory (e.g., for the GTDB database; Table 1), the accuracy remains high.

CONSULT is more effective than existing methods such as Kraken and Bowtie when the queries are phylogenetically distant from their closest match in the reference dataset. While one may hope that denser reference sets will diminish the need for such distant matching, our results on the GORG dataset demonstrate that state-of-the-art microbial datasets are far from capturing the diversity of life with the ≈ 8% distant where existing methods are accurate. Note that every genome in GORG is a single-cell assembled bacterial genome sampled randomly from the ocean; thus, these data are not exotic species put together just to challenge methods. Our results indicate that detecting even the domain of a read will require allowing many mismatches for the foreseeable future.

While we tested contamination in the context of genome skimming, we note that contamination in sequencing reads is a pervasive problem that can impact other analyses as well (86, 87, 88). It can lead to inaccurate characterization of gene content and metabolic functions (89, 90), improper inference of phylogenetic relationships (91, 92), and biases in genotype calling and population genomics (93, 94). Contamination is also known to infiltrate reference genomes stored in public databases (95) and is particularly problematic when endogenous DNA is scarce (96, 97, 98, 99). Thus, CONSULT may find applications outside the settings tested here.

We also showed that inclusion filtering of mitochondrial reads using CONSULT enabled to generate complete and accurate assemblies for very poorly preserved samples where read coverage is not sufficient to use other methods. Workflow that we developed is an example of what (100) called a “hybrid assembly method” for taking advantage of references. By searching reads against all available organelle genomes and allowing mismatches, it eliminates the bias associated with template based assembly using a single reference; at the same time, it permits flexibility of *de novo* assembly. Using CONSULT for this application is reference agnostic and thus can be utilized on mislabelled samples or samples of unknown identity. Importantly, note that our data clearly show that there is no need to have the same species or even any representative from the same genus in the reference set for the filtering to work successfully.

Finally, at its core, CONSULT is simply a read matching method. Thus, while we focused on contamination detection and organelle read detection, our algorithm can also be adopted for other applications such as metagenomic profiling, OTU picking, and any question where inexact read matching is needed. Moreover, here, performed contamination filtering using an exclusion-filter; however, a tantalizing opportunity that CONSULT may enable by allowing distant matches is inclusion filtering: find reads that seem to belong to the group of interest if assembled genomes from that phylogenetic group are available. Our results on organelle genomes, which used CONSULT as an inclusion filter, support the viability of this approach. However, inclusion filters will have to contend with contamination in the reference assemblies, which would likely require new algorithmic innovations. We leave the exploration of such applications to future work.

## Availability of data and materials

CONSULT is implemented in `C++11` with some x86 assembly code; it is (trivially) parallelized using OpenMP (101) to read the library and perform the search. The software is available publicly at https://github.com/noraracht/CONSULT. Scripts and summary data tables are publicly available on https://github.com/noraracht/lsh_scripts. Raw data used in the manuscript is deposited in https://github.com/noraracht/lsh_raw_data. The detailed description of genomic datasets used in our experiments, accession numbers of the assemblies and the exact commands used to simulate genome skims and analyze data are provided in Supplemental Material.

## Author Contributions

All authors conceived the idea. SM and VB developed a theoretical model. ER implemented the pipeline and performed experiments. All authors contributed to the analyses of data and the writing. All authors read and approved the final manuscript.

