## Supplemental information for "CONSULT: Accurate contamination removal using locality-sensitive hashing"

### A Supplementary Tables

| **Tag (bits)** | **$k$-mers count in database (billion)** | **Count of fully populated rows(%), $l$ = 0** | **Count of fully populated rows(%), $l$ = 1** |
|---|---|---|---|
| 0 | 5.838554966 | 0.354 | 0.364 |
| 2 | 6.210280798 | 0.330 | 0.339 |

Table S1: **$k$-mer counts in CONSULT database constructed with variable $t$.** Databases were built with default settings using genomes from Bacterial and Archaeal Kraken. Data demonstrate that larger tag facilitates more uniform distribution of signatures within lookup table which leads to better utilization of signature array space and ultimately allows to populate database more efficiently.

Table S2: Experiments on settings of CONSULT, namely number of hash bits ($h$), number of hash functions ($l$), number of $k$-mers per has-value ($b$), and tag size in bits ($t$). All tests were performed on TOL query set that was searched against TOL database. To avoid biases we fixed positions of the $k$-mer bits that were selected to generate signatures. For each setting, we show size of the library on the disk, recall for TOL queries searched against TOL database, and percentage of all $k$-mers among 14.026601864 billion minimized $k$-mers that fit the library. Parameter selection was mainly driven by our desire to maximize weighted recall and intention to keep database footprint under 128G.

| $h$ | $l$ | $b$ | $t$ | Disk (GB)† | Recall | % k-mer | Notes |
|---|---|---|---|---|---|---|---|
| Changing $l$ and $b$ | | | | | | | |
| 15 | 1 | 7 | 2 | 78 | 0.672 | 0.474 | Reduced recall compared to default |
| 15 | 2 | 7 | 2 | 114 | 0.696 | 0.570 | Default |
| 15 | 3 | 7 | 2 | 135 | 0.705 | 0.570 | Memory exceeds 128GB |
| 15 | 1 | 10 | 2 | 95 | 0.680 | 0.570 | Reduced recall compared to default |
| 15 | 2 | 10 | 2 | 120 | 0.700 | 0.570 | Memory exceeds 128GB† |
| 15 | 3 | 10 | 2 | 148 | 0.707 | 0.570 | Memory exceeds 128GB |
| Changing $b$ | | | | | | | |
| 15 | 2 | 4 | 2 | 76 | 0.678 | 0.394 | Reduced recall |
| 15 | 2 | 7 | 2 | 114 | 0.696 | 0.570 | Default |
| 15 | 2 | 10 | 2 | 120 | 0.700 | 0.570 | Increased memory with no benefit |
| 15 | 2 | 13 | 2 | 125 | 0.702 | 0.570 | Increased memory with no benefit |
| 15 | 2 | 16 | 2 | 128 | 0.704 | 0.570 | Increased memory with no benefit |
| Changing $h$ | | | | | | | |
| 15 | 2 | 7 | 2 | 114 | 0.696 | 0.570 | Default |
| 14 | 2 | 16 | 2 | 73 | 0.689 | 0.379 | Reduced recall and $k$-mer representation |

†: Disk space used to keep the library, as computed using the `du` command without the -apparent-size option. Note that the actual space used (in RAM) is slightly higher and is closer to what `du -apparent-size -h` would produce. For example, for the default settings, the memory usage in RAM is 120GB, whereas `du -h` produces 114GB on and `du -apparent-size -h` produces 120GB.

| Bin | Number of genomes in bin | Minimum distance encountered within bin | Maximum distance encountered within bin | Mean distance for all species within bin |
|---|---|---|---|---|
| 0 | 10 | 0.000 | 0.000 | 0.000 |
| (0 - 5] | 43 | 0.000 | 0.049 | 0.016 |
| (5 - 15] | 19 | 0.058 | 0.146 | 0.107 |
| (15 - 25] | 17 | 0.157 | 0.244 | 0.210 |
| >25 | 21 | 0.263 | 1.000 | 0.729 |

Table S3: **Bin assignment based on Mash (63) distances.**

| Species | Genome size (M) | Assembly accession | URL |
|---|---|---|---|
| *Arabidopsis thaliana* | 119.167 | GCF_000001735.4 | https://www.ncbi.nlm.nih.gov/assembly/GCF_000001735.4 |
| *Arabidopsis lyrata* | 202.97 | GCF_000004255.2 | https://www.ncbi.nlm.nih.gov/assembly/GCF_000004255.2 |
| *Carya illinoinensis* | 649.75 | GCA_011037805.1 | ftp://parrot.genomics.cn/gigadb/pub/10.5524/100001_101000/100571/Cil.genome.fa.gz |
| *Carya cathayensis* | 721.33 | GCA_011037825.1 | ftp://parrot.genomics.cn/gigadb/pub/10.5524/100001_101000/100571/Cca.genome.fa.gz |
| *Nicotiana sylvestris* | 2221.99 | GCF_000393655.1 | https://www.ncbi.nlm.nih.gov/assembly/GCF_000393655.1 |
| *Zea mays* | 2182.61 | GCF_000005005.2 | https://www.ncbi.nlm.nih.gov/assembly/GCF_000005005.2 |
| *Oryza sativa* | 382.63 | GCF_001433935.1 | https://www.ncbi.nlm.nih.gov/assembly/GCF_001433935.1 |
| *Coffee arabica* | 1094.45 | GCF_003713225.1 | https://www.ncbi.nlm.nih.gov/assembly/GCF_003713225.1 |
| *Prunus persica* | 212.77 | GCF_000346465.2 | https://www.ncbi.nlm.nih.gov/assembly/GCF_000346465.2 |
| *Bathycoccus prasinos* | 15.07 | GCF_002220235.1 | https://www.ncbi.nlm.nih.gov/assembly/GCF_002220235.1 |

Table S4: **List of plant species added to TOL query set.**

| Species | Run | URL |
|---|---|---|
| *Drosophila bipectinata* | SRR6425989 | https://www.ncbi.nlm.nih.gov/sra/?term=SRR6425989 |
| *Drosophila erecta* | SRR6425990 | https://www.ncbi.nlm.nih.gov/sra/?term=SRR6425990 |
| *Drosophila ananassae* | SRR6425991 | https://www.ncbi.nlm.nih.gov/sra/?term=SRR6425991 |
| *Drosophila biarmipes* | SRR6425992 | https://www.ncbi.nlm.nih.gov/sra/?term=SRR6425992 |
| *Drosophila mauritiana* | SRR6425993 | https://www.ncbi.nlm.nih.gov/sra/?term=SRR6425993 |
| *Drosophila eugracilis* | SRR6425995 | https://www.ncbi.nlm.nih.gov/sra/?term=SRR6425995 |
| *Drosophila mojavensis* | SRR6425997 | https://www.ncbi.nlm.nih.gov/sra/?term=SRR6425997 |
| *Drosophila persimilis* | SRR6425998 | https://www.ncbi.nlm.nih.gov/sra/?term=SRR6425998 |
| *Drosophila simulans* | SRR6425999 | https://www.ncbi.nlm.nih.gov/sra/?term=SRR6425999 |
| *Drosophila virilis* | SRR6426000 | https://www.ncbi.nlm.nih.gov/sra/?term=SRR6426000 |
| *Drosophila pseudoobscura* | SRR6426001 | https://www.ncbi.nlm.nih.gov/sra/?term=SRR6426001 |
| *Drosophila sechellia* | SRR6426002 | https://www.ncbi.nlm.nih.gov/sra/?term=SRR6426002 |
| *Drosophila willistoni* | SRR6426003 | https://www.ncbi.nlm.nih.gov/sra/?term=SRR6426003 |
| *Drosophila yakuba* | SRR6426004 | https://www.ncbi.nlm.nih.gov/sra/?term=SRR6426004 |

Table S5: **List of SRRs and URLs for Drosophila species used in real data experiment.**

| Run | Species | Contig length(bp) | mtDNA(%) | GenBank ID | Available genus | Available species |
|---|---|---|---|---|---|---|
| | | | Unassembled samples | | | |
| SRR12432370 | *Apodemus sylvaticus* | N/A | N/A | N/A | 6 | 1 |
| SRR12432443 | *Argyropelecus olfersii* | N/A | N/A | N/A | 0 | 0 |
| SRR12432479 | *Branta leucopsis* | N/A | N/A | N/A | 1 | 0 |
| SRR12432364 | *Chelon ramada* | N/A | N/A | N/A | 1 | 0 |
| SRR12432468 | *Corvus cornix* | N/A | N/A | N/A | 12 | 1 |
| SRR12432397 | *Cygnus olor* | N/A | N/A | N/A | 4 | 1 |
| SRR12432591 | *Emberiza citrinella* | N/A | N/A | N/A | 12 | 0 |
| SRR12432369 | *Eptesicus serotinus* | N/A | N/A | N/A | 0 | 0 |
| SRR12432332 | *Gallinago gallinago* | N/A | N/A | N/A | 1 | 0 |
| SRR12432390 | *Ichthyosaura alpestris* | N/A | N/A | N/A | 0 | 0 |
| SRR12432365 | *Maurolicus muelleri* | N/A | N/A | N/A | 0 | 0 |
| SRR12432368 | *Megaptera novaeangliae* | N/A | N/A | N/A | 0 | 1 |
| SRR12432371 | *Microtus arvalis* | N/A | N/A | N/A | 7 | 1 |
| SRR12432366 | *Myotis mystacinus* | N/A | N/A | N/A | 29 | 0 |
| SRR12432391 | *Pipistrellus pygmaeus* | N/A | N/A | N/A | 2 | 0 |
| SRR12432352 | *Raniceps raninus* | N/A | N/A | N/A | 0 | 0 |
| SRR12432355 | *Sicista betulina* | N/A | N/A | N/A | 1 | 0 |
| SRR12432363 | *Trachipterus arcticus* | N/A | N/A | N/A | 1 | 0 |
| | | | Poor quality assemblies | | | |
| SRR11679515 | *Tamias sibiricus* | 3746 | 0.0049 | MT410867 | 7 | 1 |
| SRR11679527 | *Lullula arborea* | 3018 | 0.0114 | MT410892 | 0 | 0 |
| SRR11679474 | *Callionymus reticulatus* | 4644 | 0.0138 | MT410925 | 2 | 0 |
| SRR11537214 | *Asio flammeus* | 10466 | 0.0279 | MN122899 | 1 | 1 |
| SRR11537185 | *Calidris alpina* | 9562 | 0.0393 | MN122893 | 3 | 0 |
| SRR11537153 | *Lycodes vahlii* | 8147 | 0.2552 | MT410895 | 5 | 0 |
| | | | Good quality assemblies (positive control) | | | |
| SRR11679529 | *Luscinia luscinia* | 14276 | 0.0164 | MT410894 | 2 | 0 |
| SRR11679531 | *Phasianus colchicus* | 16773 | 0.0219 | MT410882 | 1 | 1 |
| SRR11537201 | *Merlangius merlangus* | 16660 | 0.0298 | MN122861 | 0 | 1 |
| SRR11537177 | *Merops apiaster* | 13884 | 0.0406 | MN122929 | 1 | 0 |
| SRR11679575 | *Oncorhynchus mykiss* | 16759 | 0.0820 | MT410879 | 11 | 1 |
| SRR11537188 | *Buteo buteo* | 13998 | 0.1356 | MN122916 | 2 | 1 |
| SRR11679510 | *Physeter catodon* | 16491 | 0.3210 | MT410874 | 0 | 1 |
| SRR11679507 | *Taurulus bubalis* | 16854 | 0.4018 | MT410868 | 0 | 0 |
| SRR11679528 | *Arvicola amphibius* | 16356 | 0.5788 | MN122828 | 0 | 0 |
| SRR11679539 | *Sciurus vulgaris* | 16511 | 0.8176 | MN122875 | 11 | 1 |
| SRR11537162 | *Arnoglossus laterna* | 15958 | 1.0269 | MN122822 | 2 | 0 |
| SRR11679545 | *Vipera berus* | 12733 | 1.2580 | MN122848 | 0 | 1 |
| SRR11537150 | *Centrolabrus exoletus* | 16494 | 1.6414 | MT410926 | 0 | 0 |
| SRR11679513 | *Lagenorhynchus albirostris* | 16393 | 2.0310 | MT410901 | 2 | 1 |
| SRR11679516 | *Delphinus delphis* | 16386 | 2.8277 | MT410915 | 1 | 1 |
| SRR11537206 | *Vipera berus* | 12733 | 4.0683 | MN122824 | 0 | 1 |
| SRR11537189 | *Milvus milvus* | 17883 | 4.7909 | MN122837 | 1 | 0 |
| SRR11679555 | *Chroicocephalus ridibundus* | 16768 | 5.6225 | MN122820 | 1 | 1 |

Table S6: **Samples from Denmark sequencing project (75) used in mitochondrial assembly experiment.** Unassembled subset (n=18) represents species that failed mitochondrial assembly in Denmark study. Poor quality group (n=6) was assembled with short contigs of 3–10.5 kbp. Good quality assemblies (n=18) were randomly selected samples with relatively long contig length. Genus count and species count represent count of samples of the same genus or species present in RefSeq release used to construct mitochondrial CONSULT library.

| Genome | Assembly accession | Bioproject | Species |
|---|---|---|---|
| G000019605 | GCF_000019605.1 | PRJNA224116 | *Candidatus Korarchaeum cryptofilum* |
| G000231015 | GCF_000231015.2 | PRJNA224116 | *Desulfurococcus fermentans* |
| G000816105 | GCF_000816105.1 | PRJNA224116 | *Thermococcus guaymasensis* |
| G001510295 | GCA_001510295.1 | PRJNA279271 | *Thaumarchaeota archaeon* |
| G001674955 | GCA_001674955.1 | PRJNA289040 | *Candidatus Nitrosopumilus sp.* |
| G000007185 | GCA_000007185.1 | PRJNA294 | *Methanopyrus kandleri* |
| G000204585 | GCA_000204585.1 | PRJNA52465 | *Candidatus Nitrosoarchaeum limnia* |
| G000421185 | GCF_000421185.1 | PRJNA224116 | *Ornithinimicrobium pekingense* |
| G001315825 | GCA_001315825.1 | PRJDB782 | *Metallosphaera hakonensis* |
| G001577775 | GCF_001577775.1 | PRJNA224116 | *Pyrococcus kukulkanii* |
| G001940645 | GCA_001940645.1 | PRJNA319486 | *Candidatus Heimdallarchaeota archaeon* |
| G000011125 | GCA_000011125.1 | PRJNA211 | *Aeropyrum pernix* |
| G000221185 | GCF_000221185.1 | PRJNA224116 | *Thermococcus sp.* |
| G000770635 | GCF_000770635.1 | PRJNA224116 | *Pontibacillus yanchengensis* |
| G001316265 | GCF_001316265.1 | PRJNA224116 | *Vulcanisaeta sp.* |
| G001628455 | GCA_001628455.1 | PRJNA289734 | *Marine group II euryarchaeote* |
| G001940655 | GCA_001940655.1 | PRJNA288027 | *Candidatus Lokiarchaeota archaeon* |
| G000022365 | GCF_000022365.1 | PRJNA224116 | *Thermococcus gammatolerans* |
| G000307305 | GCF_000307305.1 | PRJNA224116 | *Enterobacteriaceae bacterium* |
| G000955905 | GCF_000955905.1 | PRJNA224116 | *Candidatus Nitrosotenuis cloacae* |
| G001515215 | GCA_001515215.1 | PRJNA298487 | *Hadesarchaea archaeon* |
| G001679155 | GCF_001679155.1 | PRJNA224116 | *Moraxella nonliquefaciens* |
| G000145295 | GCF_000145295.1 | PRJNA224116 | *Methanothermobacter marburgensis* |
| G000375685 | GCA_000375685.1 | PRJNA165545 | *crenarchaeote SCGC* |
| G001189275 | GCA_001189275.1 | PRJNA258558 | *Pyrobaculum sp.* |
| G001560165 | GCF_001560165.1 | PRJNA224116 | *Sulfolobus acidocaldarius* |
| G001919175 | GCA_001919175.1 | PRJNA297196 | *Crenarchaeota archaeon* |
| Cca.genome.fa.gz | GCA_011037825.1 | PRJNA435846 | *Carya cathayensis* |
| Cil.genome.fa.gz | GCA_011037805.1 | PRJNA435846 | *Carya illinoinensis* |
| TAIR10.1 | GCF_000001735.4 | PRJNA10719 | *Arabidopsis thaliana* |

Table S7: **Subset of TOL samples used in running time analysis.** Samples were selected randomly.

| **Run** | **Bioproject** | **Species** |
|---|---|---|
| SRR6425989 | PRJNA427774 | *Drosophila bipectinata* |
| SRR6425990 | PRJNA427774 | *Drosophila erecta* |
| SRR6425991 | PRJNA427774 | *Drosophila ananassae* |
| SRR6425992 | PRJNA427774 | *Drosophila biarmipes* |
| SRR6425993 | PRJNA427774 | *Drosophila mauritiana* |
| SRR6425995 | PRJNA427774 | *Drosophila eugracilis* |
| SRR6425997 | PRJNA427774 | *Drosophila mojavensis* |
| SRR6425998 | PRJNA427774 | *Drosophila persimilis* |
| SRR6425999 | PRJNA427774 | *Drosophila simulans* |
| SRR6426000 | PRJNA427774 | *Drosophila virilis* |
| SRR6426001 | PRJNA427774 | *Drosophila pseudoobscura* |
| SRR6426002 | PRJNA427774 | *Drosophila sechellia* |
| SRR6426003 | PRJNA427774 | *Drosophila willistoni* |
| SRR6426004 | PRJNA427774 | *Drosophila yakuba* |
| SRR12129012 | PRJNA643549 | *Drosophila melanogaster* |

Table S8: **Drosophila species used as query sequences in running time analysis.**

### B Supplementary Figures

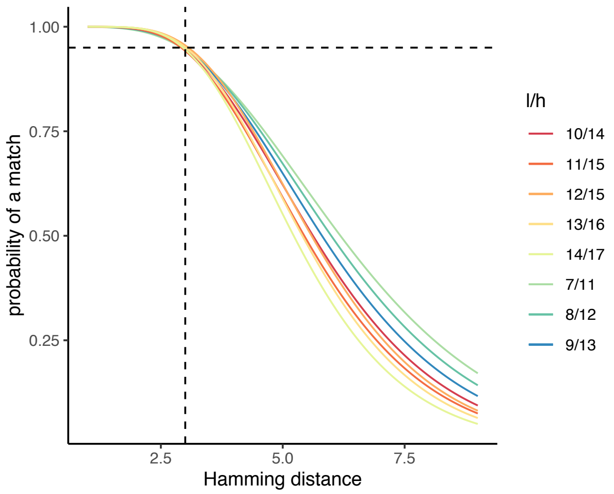


(a)

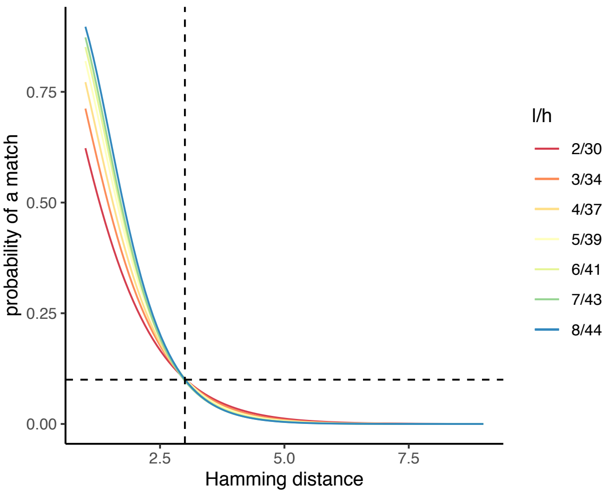


(b)

Figure S1: **Controlling true positive rate.** Assuming we want a fraction $\rho$ of $k$-mers in a read to match to sequences with $p$ hamming distance to the read, we need $l$ hashes and $h = \frac{\log(1-(1-\rho)^{\frac{1}{l}})}{\log(1-\frac{p}{k})}$ bits per hash to achieve the desired level. Figures show probability of a 32-mers matching another 32-mer ($p(d)$) if their distance $d$ is the value shown on x-axis, for various choices of $l$, setting $h$ such that $\rho(3) = 0.10$ (on the right) or $\rho(3) = 0.95$ (on the left) ($p = 3$ and $\rho(3)$ shown with dotted lines).

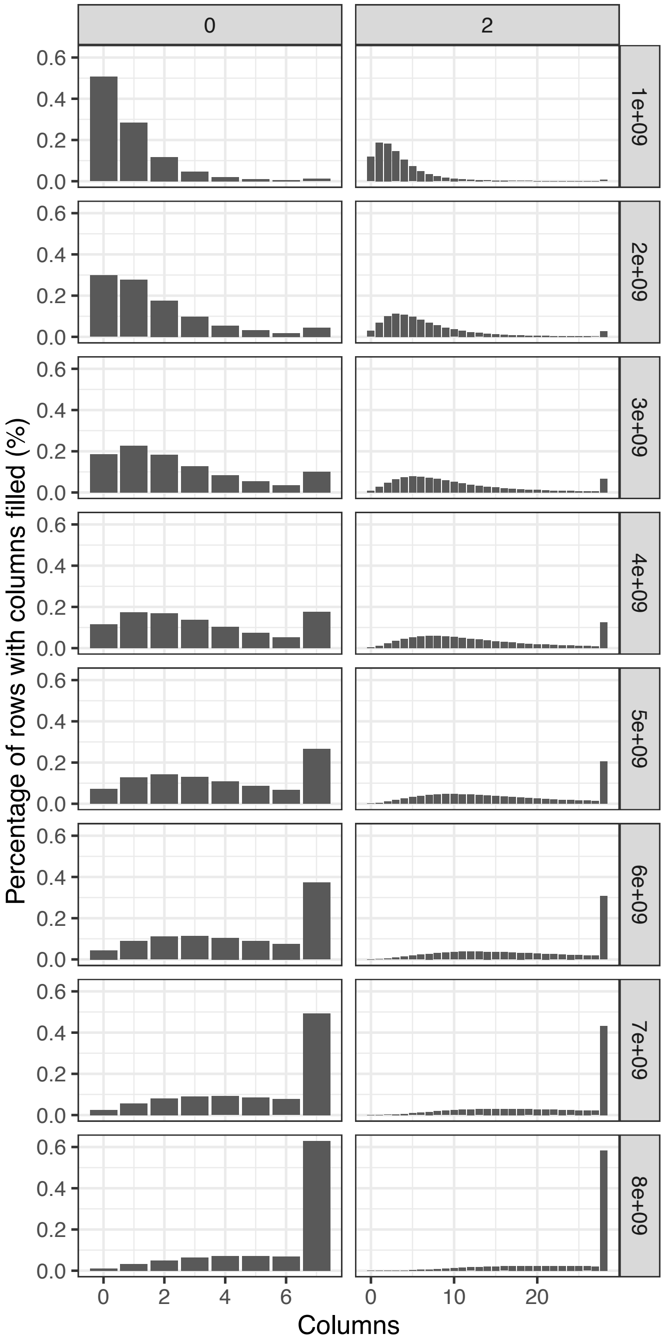


Figure S2: **Dynamics of filling out lookup tables during database construction.** Each bar shows the percentage of rows where the x-axis is the number of columns that are populated with pointers to the encoding table. Row panels show counts after a number of $k$-mers were added to the database while column panels show tag size (0 or 2). Experiment represents a construction of TOL database. With tag 0, many rows starts to fill up around 3 billion $k$-mers, showing that row usage is not uniform. For most levels between 1 billion and 6 billion $k$-mers, fewer rows are full with tag 2 than tag 0. This pattern is the motivation for using $t = 2$ as the default.

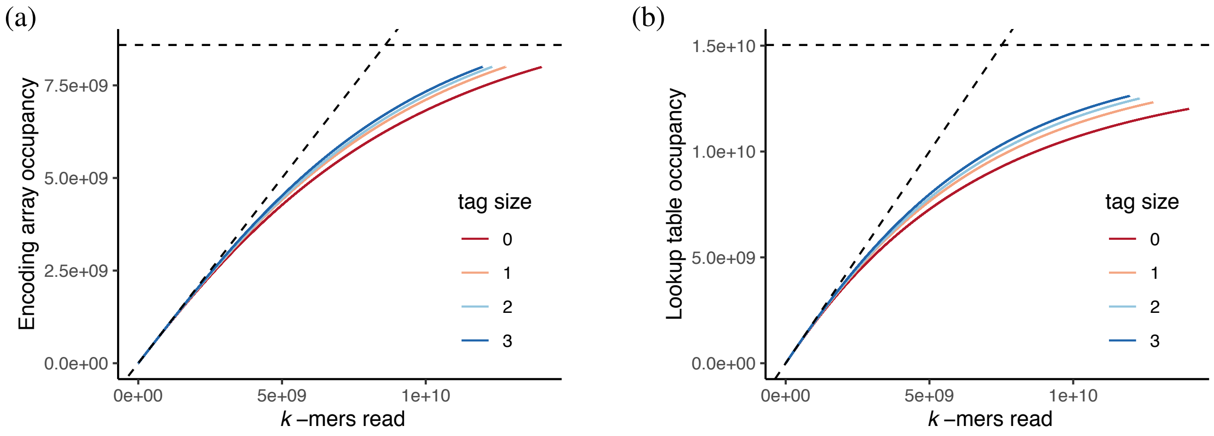


Figure S3: **Effect of tag size on efficiency of $k$-mer inclusion during database construction.** x-axis shows the number of $k$-mers that were read during database construction at given point of time. y-axis represents (a) total number of encodings that were added to encoding array, and (b) total number of signatures that were added to lookup table. Experiment indicates that with $t = 0$, we have less efficient utilization of map capacity. Experiment was performed on TOL database constructed with default settings and $t \in \{0, 1, 2, 3\}$.

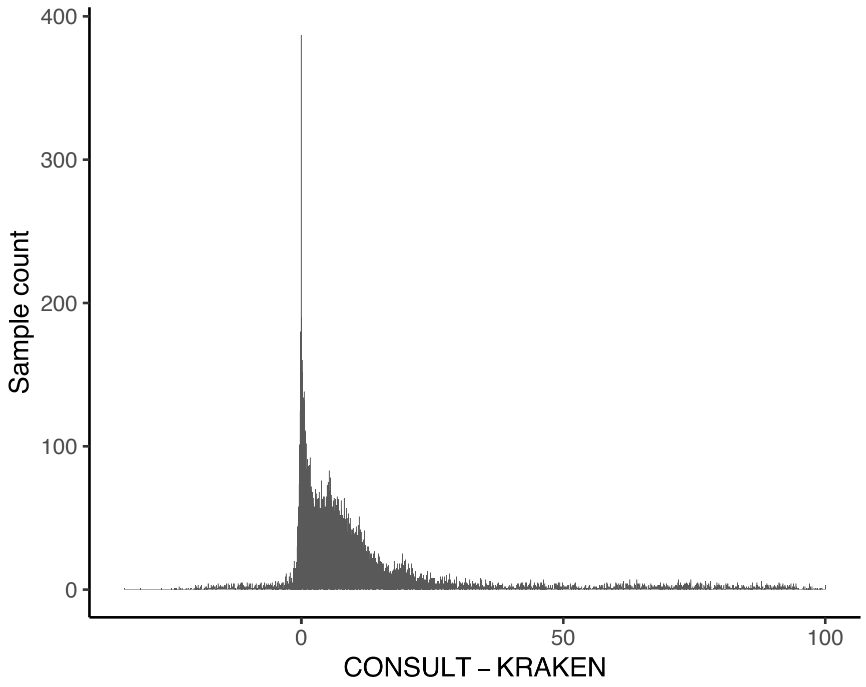


Figure S4: **Difference in percentage of classified reads between CONSULT and Kraken.** The x-axis shows the difference between the percentage of reads in each genome that are classified using CONSULT and Kraken. Thus, positive difference indicates that CONSULT classifies more reads for a given sample than Kraken. The histogram is shown over all $> 12,000$ GORG samples queried against the GTDB reference library with default settings (bin size=0.1%).

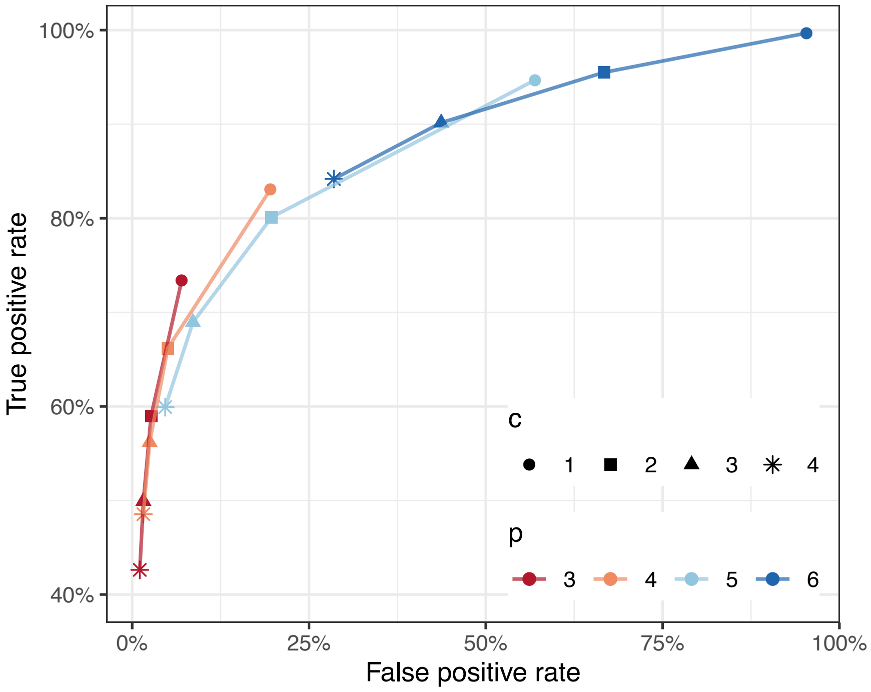


Figure S5: **Parameter exploration of CONSULT.** Performance of CONSULT with GORG samples queried against GTDB at variable $p \in \{3, 4, 5, 6\}$ and $c \in \{1, 2, 3, 4\}$. CONSULT GTDB library was constructed with default settings.

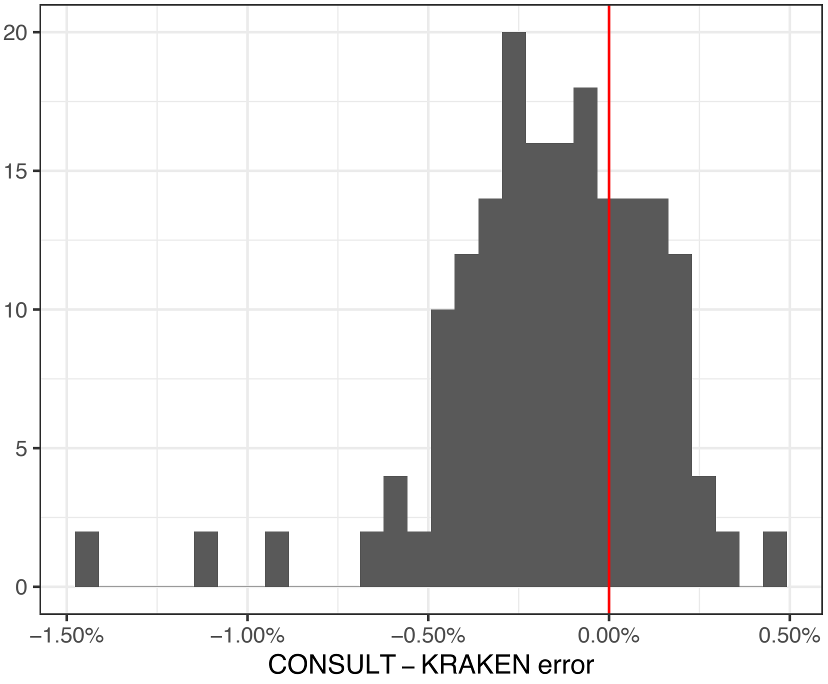


Figure S6: **Difference in relative error in distance estimates computed after query filtering performed with CONSULT or Kraken.** Analysis was done on Drosophila sequences that where searched against GTDB. We used default ($p = 3, c = 1$) run parameters for CONSULT and 0.0 confidence level for Kraken. Distances were computed using Skmer. Samples on a positive side have higher error with CONSULT. Samples on a negative side have higher error with Kraken. y-axis represents sample count.

### C Supplementary method details and commands

Here we provide the exact procedures and commands that we used to run external software throughout our experiments.

#### Genome skim simulation

We simulated short reads with length $l$ = 150 and coverage $c$, in single read mode with default error and quality profiles of Illumina HiSeq 2500 using ART version 2.5.8 with command

```
art_illumina -ss HS25 -i FASTA_FILE -l 150 -f c -na -s 10 -o FASTQ_FILE
```

#### Downsampling reads

To subsample reads down a specified number of reads $n$ we used seqtk version 1.3r106 with command

```
seqtk sample -s150 INPUT_FASTQ_FILE n > OUTPUT_FASTQ_FILE
```

#### Kraken reference library construction

In this study we used Kraken version 2.

- To construct standard Kraken reference library we used default command

  ```
  kraken2-build --standard --no-masking --use-ftp --db DATABASE_NAME
  ```

- To build custom Kraken reference library we used a set of commands below:

  1. Download taxonomy

  ```
  kraken2-build --download-taxonomy --no-masking --use-ftp
  --db DATABASE_NAME
  ```

  2. Rename file extensions to .fa

  ```
  find .  -name "*.fna" -exec sh -c 'mv "$1" "${1%.fna}.fa"' _ {} \;
  ```

  3. Add custom genomes to the reference library

  ```
  find genomes/ -name '*.fa' -print0 | xargs -0 -I -n1 kraken2-build
  --no-masking --add-to-library {} --db DATABASE_NAME
  ```

  4. To build database with specified $k$-mer length $k$, minimizer length $l$ and number of wind-carding positions $s$ we used

  ```
  kraken2-build --build --no-masking --kmer-len k --minimizer-len l
  --minimizer-spaces s --use-ftp --db DATABASE_NAME
  ```

#### Kraken reference library querying

To query Kraken reference library at variable confidence level $\alpha$ we used

```
kraken2 --use-names --threads 24 --report REPORT_FILE_NAME
--db DATABASE_NAME --confidence α --classified-out CLASSIFIED_FASTQ_FILE
--unclassified-out UNCLASSIFIED_FASTQ_FILE QUERY_FASTQ_FILE >
KRAKEN_OUTPUT_FILE
```

#### CLARK(-S) reference library construction and querying

We used CLARK version 1.2.6.1.

During database construction, CLARK was able to set target IDs for 9976 genomes out of 10470 sample that we aimed to include. Remaining TOL genomes were added manually by setting taxonomic rank for a given sample to Proteobacteria phylum and specifying taxonomy ID as 1224.

- Custom CLARK database of discriminative $k$-mers was built in `<DIR_DB/>` the following way:

  1. We created the directory "Custom" inside <DIR_DB/>

  2. We copied the sequences of interest (in our case reference TOL fasta files with accession numbers) in the "Custom"

  3. We ran

  ```
  ./set_target.ssh <DIR_DB/> custom --phylum
  ```

  We note we set taxonomy rank to phylum. The default taxonomy rank is species.

- CLARK database was queried at default mode of classification (i.e., "-m 1") with script

  ```
  ./classify_metagenome.sh -O SAMPLE_FASTQ_FILE -R RESULT_CSV -n 24
  ```

- CLARK-S databases of discriminative **spaced** $k$-mers was created on top of custom CLARK database using script

  ```
  ./buildSpacedDB.sh
  ```

- CLARK-S classification was ran with "full" mode ("-m 0" identifier) using script

  ```
  ./classify_metagenome.sh -O SAMPLE_FASTQ_FILE -R RESULT_CSV -n 24
  --spaced -m 0
  ```

#### Bowtie2 reference index construction and alignment

We used Bowtie2 version 2.4.1.

- To index reference genomes we used command

  ```
  bowtie2-build <reference_in> <bt2_base> -threads 24
  ```

  where <reference_in> refers to reference sequences (in our case genomes from TOL dataset) that were concatenated into single fasta file prior to index construction and <bt2_base> is the basename of the index files to write.

- To run alignment we used

  ```
  bowtie2 -x <bt2_base> -U INPUT_FASTQ_FILE --local
  --very-sensitive -p 24 -S RESULT_SAM_FILE
  ```

### Manipulation and post-processing of read alignments

We used SAMtools version 1.9.

- To obtain alignment statistics

  ```
  samtools stats INPUT_ALN_SAM > OUTPUT_STATS_TXT
  samtools flagstat INPUT_ALN_SAM > OUTPUT_STATS_TXT
  ```

### CONSULT reference library construction and querying

- To build CONSULT reference libraries we used version 17.1 of mapping software

  1. We compiled script using
     ```
     g++ main_map.cpp -std=c++11 -O3 -o main_map
     ```
  2. To construct reference database with default settings we ran
     ```
     ./main_map -i INPUT_FASTA_FILE -o DB_FOLDER_NAME
     ```

  To avoid biases during testing we fixed positions of bits that are selected during signature generation. In released version positions are selected randomly.

  For $h$=15 we used $l$ = 0 $bits \in \{30, 28, 25, 24, 22, 21, 20, 17, 14, 13, 11, 5, 3, 2, 1\}$

  $l$ = 1 $bits \in \{30, 29, 26, 24, 20, 19, 16, 14, 13, 12, 10, 7, 3, 1, 0\}$

  $l$ = 2 $bits \in \{30, 29, 26, 23, 21, 20, 19, 18, 15, 11, 9, 7, 5, 3, 2\}$

- CONSULT database was queried using version 17.4 of search software

  1. To compile
     ```
     g++ main_search.cpp -std=c++11 -fopenmp -O3 -o main_search
     ```
  2. To query sequence reads against reference database we ran
     ```
     ./main_search -i DB_FOLDER_NAME -c 1 -t 24 -q QUERY_FOLDER_NAME
     ```

     **where arguments are:**

     -i - name of the reference database

     -c - the lowest number of $k$-mers that is required to mark sequencing read as classified. For instance, if at least one $k$-mer match is enough to classify a read, "c" should be set to 1. If at least two $k$-mer matches are required to call read a match, "c" should be set to 2.

     -t - number of threads

     -q - name of the folder where queries are located

  For measuring running time of CONSULT, we split query file using Linux `split -n l/48` and provided a folder containing all the files to the tool; we added split time to the total running time.

#### Generation of $k$-mer sets

To estimate $k$-mer frequencies we used Jellyfish version 2.3.0.

- To compute 35bp and 32bp canonical $k$-mer profiles of fasta genomic references we used

```
jellyfish count -m 35 -s 100M -t 24 -C INPUT_FASTA_FILE -o COUNT_JF_FILE
jellyfish count -m 32 -s 100M -t 24 -C INPUT_FASTA_FILE -o COUNT_JF_FILE
```

- To output a list of all the $k$-mers in the file associated with their counts

```
jellyfish dump COUNT_JF_FILE > OUTPUT_FASTA_FILE
```

#### $k$-mer minimization

Minimization was performed using custom c++ minimization script version 3.0 which is available for public use. This script accepts as an input Jellyfish fasta file containing 35 bp canonical $k$-mers extracted from reference genomes and outputs their 32 bp minimizers in fasta format.

- To compile the script we used

```
g++ minimization_v3.0.cpp -std=c++11 -o main_minimization
```

- We ran minimization with the command

```
./main_minimization -i INPUT_FASTA_FILE -o MIN_OUTPUT_FASTA_FILE
```

#### Computation of genomic distances

To estimate genomic distances we used Mash 1.1 and Skmer 3.0.2.

- To compute genomic distance with Mash we used

```
mash dist FASTQ_FILE_ONE FASTQ_FILE_TWO
```

- To compute genomic distance with Skmer we used

```
skmer reference FASTQ_DIRECTORY -p 24 -o REF_DISTANCE_MATRIX
```

#### Preprocessing of Drosophila sequencing files

We used BBTools version 38.59 to preprocess real data sequencing reads.

- To decontaminate .fastq files we used

```
bbduk.sh in1=FASTQ_READ1 in2=FASTQ_READ2 out1=FASTQ_READ1
out2=FASTQ_READ2 ref=adapters,phix ktrim=r k=23 mink=11 hdist=1 tpe tbo
```

- To deduplicate reads we used

```
dedupe.sh in1=FASTQ_READ1 in2=FASTQ_READ2 out=DEDUP_OUTPUT_FASTQ_FILE
```

- To reformat deduplicated output files we used

```
reformat.sh in=DEDUP_OUTPUT_FASTQ_FILE out1=FASTQ_READ1 out2=FASTQ_READ2
```

- To merge paired-end reads .fastq we used

```
bbmerge.sh in1=FASTQ_READ1 in2=FASTQ_READ2 out1=OUTPUT_FASTQ_FILE
```

#### Preprocessing of mitochondrial sequencing files

- We used AdapterRemoval version 2.3.1 to remove adapters from raw sequencing data

```
AdapterRemoval -file1 FASTQ_READ1 -file2 FASTQ_READ2 -basename FASTQ_trmd
```

- We used BBTools version 38.59 to merge paired-end reads

```
bbmerge.sh in1=FASTQ_READ1 in2=FASTQ_READ2 out=merged.fastq
outu1=read1_unmerged.fastq outu2=read2_unmerged.fastq
```

- Subsequently we concatenated merged and unmerged outputs

```
cat merged.fastq read1_unmerged.fastq read2_unmerged.fastq > OUTPUT_FILE
```

#### To collect classified reads from CONSULT we used

- We used BBTools version 38.59

```
filterbyname.sh in=INIT_FILE names=UCSEQ_FILE out=CSEQ_FILE include=f
int=f overwrite=true
```

#### Assembling mitochondrial reads

To assemble mitochondrail reads we used SPAdes genome assembler version 3.15.0.

- On unfiltered reads we ran

```
spades.py -plasmid -only-assembler -s INPUT_FASTQ -t 24 -m 120
-o OUTPUT_DIR
```

- To assemble filtered reads we ran

```
spades.py -only-assembler -s INPUT_FASTQ -t 24 -m 120 -o OUTPUT_DIR
```

#### Mitochondrial annotation

Annotation of mitochondrial assemblies was done with MITOS version 2.0.8 using RefSeq89 Metazoa reference and vertebrate mitochondrial code 2.

- We ran

```
runmitos.py -i INPUT_FASTA -c 2 -o OUTPUT_DIR -linear -r REFSEQVER -R
REFDIR
```

- To confirm identity of largest mitochondrial contigs for unfiltered assemblies we used MitoZ version 2.4

```
python MitoZ.py annotate -genetic_code auto -clade Chordata -outprefix
test -thread_number 8 -fastafile INPUT_FASTA
```

**Algorithm 2** CONSULT algorithm. Notations: $\mathcal{S}$: all reference sequences. Defaults: $m = 35, k = 32, h = 15, l = 2, t = 2, b = 7, p = 3, c = 1, g = 33$. $[a]$ denotes $\{0, \ldots, a-1\}$. $I(s)$ returns $2h - t$ least significant bits of $s$ and $T(s)$ returns the rest. `SHIFTS` returns the number of runs of 0s and 1s in its input as tuples. `SHLD` is the x86 instruction of the same name (Double Precision Shift Left, or extended shift). Details are omitted.

```
procedure BuildLibrary(𝒮)
    E ← array of ≤ 2^g elements, each 2k bits
    for each i ∈ [l] do
        M_i ← h unique random numbers in [k − 1]
        S_i ← 2^(2h−t) × (2^t b) array of g-bit elements
        T_i ← 2^(2h−t) × (2^t) array of 1-byte elements
        S'_i ← 2^(2h−t) × (2^t b) array of (t,g)-bit tuples
    𝒦 ← Minimize({all m-mers of 𝒮}, k)
    j ← −1
    for each k-mer a ∈ 𝒦 do
        e ←LeftRightEncode(a)
        Included ← False
        for each i ∈ [l] do
            s ← Signature(M_i, e)
            if S'_i[I(s)] is not full then
                if not Included then
                    j ← j + 1
                    E[j] ← e
                    Included ← True
                Append (T(s), j) to S'_i[I(s)]
    for each row s of each S'_i do
        sort values of s, note boundaries of tags
        save 2nd elements of s to S_i[I(s)]
        save boundaries of tags in T_i[I(s)]
    save DB = (E, M, S, T) to disk
procedure Minimize(𝒦', k)                              ▷ Minimization
    ℛ ← ∅
    for each a in 𝒦' do
        Append min{all k-mers of a} to ℛ
    return ℛ ordered pseudo-randomly
procedure LeftRightEncode(a)                           ▷ Encoding
    R ← 2k-bit zeros
    for letter a_i in a do
        R_i = 1 if a_i ∈ {G, T}
        R_{i+32} = 1 if a_i ∈ {C, T}
    return R
procedure Signature(M, e)                ▷ Extract signature from e
    r ← 2h-bit zeros
    for (sh_skip, sh_keep) ∈ Shifts(M) do
        e ← Shift e left by sh_skip
        r ←SHLD (sh_keep, e, r)
    return r
procedure QueryRead(r, DB)
    l ← 0
    for k-mer a in r and its reverse complement do
        e ←LeftRightEncode(a)
        for each i ∈ [l] do
            s ← Signature(M_i, e)
            for T_i[T(s)] ≤ f < [T_i[T(s) + 1]] do
                if HD(E[S_i[I(s)][f]], e)≤ p then
                    l ← l + 1
                    if l ≥ c then
                        return r is a match
    return e is not a match
procedure HD(a,b)                                ▷ Hamming Distance
    z_up, z_low ← lower and upper k-bits of a ⊕ b
    return popcount(z_up ∨ z_low)
```
